# Detuning of the Ribosome Conformational Landscape Promotes Antibiotic Resistance and Collateral Sensitivity

**DOI:** 10.1101/2023.06.13.544509

**Authors:** Pablo Mesa, Alicia Jiménez-Fernández, Ruggero La Rosa, Rocio Espinosa, Helle Krogh Johansen, Søren Molin, Guillermo Montoya

## Abstract

Around 50% of the current antibiotic arsenal targets the ribosome, thus resistance to ribosome-targeting antibiotics poses severe challenges to antimicrobial treatments. Here, we characterize a 12-nucleotide deletion in the *rplF* gene encoding the uL6 ribosomal protein, which was identified in a tobramycin-resistant strain of *Pseudomonas aeruginosa* isolated from a cystic fibrosis patient. To understand this resistance, we determined 87 cryo-EM structures of wild-type and mutant ribosomes characterizing their conformational landscape. Our analysis reveals how detuning of the ribosome dynamics alters its rotational movement circumventing tobramycin inhibition. The mutation compromises the 50S assembly, triggering structural instability and inducing a different rotational dynamic of the 70S. We found 4 new binding sites of tobramycin, one of them exclusive of the mutant, where the binding of the antibiotic acts as an allosteric activator skipping inhibition. Our data also illustrate how chloramphenicol stabilizes the mutant ribosome, enhancing inhibition and thereby leading to collateral sensitivity.

## INTRODUCTION

During the last decades, antibiotic-resistant bacteria have become an important risk to human health ^1^. The increase in bacterial resistance together with the slow pace of discovery of new antibiotics is raising serious concerns regarding a return to the pre-antibiotic era. Almost half of the current antimicrobials target the ribosome ^2^. This large ribonucleoprotein complex of about 2.5 MDa is composed of a small subunit (30S), consisting of the 16S rRNA molecule and 20 proteins, and a large subunit (50S) that contains the 23S and 5S rRNAs and 34 proteins ^3^. Among the different families of antibiotics targeting the bacterial ribosome, aminoglycosides are widely used in the clinic ^4^. These antibiotics are frequently used for the treatment of many Gram-negative and some Gram-positive infections, and they are routinely applied against *Pseudomonas aeruginosa* infections in cystic fibrosis (CF) patients ^5^. Aminoglycosides bind near the decoding center (DC) in the 30S subunit, where mRNA and tRNA meet to translate nucleic acids to proteins ^6^. During the initiation of translation and the elongation stages of the ribosome cycle, different co-factors dock on the ribosome to support the codon-anticodon match, which triggers hydrolysis of GTP in the G-domain of ternary complexes composed by the initiation (IF) and elongation factors (EF-Tu or EF-G), GTP and aminoacyl-t-RNA^7^ ^8^. Although secondary binding sites have been reported for some members of the aminoglycoside family, e.g., neomycin has been shown to bind to helix 69 (H69) of the 23S rRNA ^9, 10^, they seem to preferentially bind to helix 44 (h44) of the 16S rRNA at clinically relevant concentrations. The aminoglycoside binding displaces the universally conserved A1492 and A1493 in the 16S rRNA ^4, 6, 11^, affecting codon-anticodon recognition in the DC, as it has been recently observed for paromomycin at 2.04 Å resolution ^12^. Consequently, the binding alters tRNA translocation, reduces translational fidelity, affects conformational changes of the ribosomal subunits interfering with the formation of inter-subunit bridges, inhibits ribosome recycling, and prevents protein synthesis ^10, 13–16^.

Resistance to aminoglycosides can be obtained by different mechanisms, including reduced uptake of the drug ^17^, chemical modification and inactivation of the antibiotic ^18^, or nucleotide modification by methyltransferases (16S-RMTases) in the antibiotic target site ^19, 20^. In our collection of more than 500 clinical isolates of *P. aeruginosa* from young CF patients, almost 20,000 mutations were identified by whole genome sequencing, of which 46 were in ribosomal protein genes ^21, 22^. Little is known about their effect on antibiotic resistance, and so far, only a few have been associated with antibiotic resistance ^23, 24^. Importantly, genetic changes in the ribosomal machinery often compromise the host cells, leading to decreased fitness when the antibiotic selective pressure fades out. Resistance development is frequently associated with enhanced sensitivity towards other antibiotics, recently described as ‘collateral sensitivity’ ^25, 26^. This phenomenon, observed in several bacterial species, suggests that the mechanistic understanding of the processes conferring collateral sensitivity may be exploited to mitigate antibiotic resistance and to improve the elimination of pathogens during infections ^27–29^. Even though strategies for the rational use of antibiotics have been proposed based on collateral sensitivity data ^30^, the underlying molecular mechanisms are still largely uncharacterized.

Here we report the functional and structural characterization of a 12-nucleotide deletion in the *rplF* gene encoding the uL6 protein of the 50S subunit. This mutation was identified in a CF clinical isolate of *P. aeruginosa* ^23^. After genetic transfer of the *rplF* mutation to the genome of the reference *P. aeruginosa* strain (PAO1), the mutant strain harboring the uL6 deletion displayed high-level resistance to a broad range of aminoglycoside antibiotics, collateral sensitivity to chloramphenicol, and severely reduced growth rate (Fig. 1). To understand how antibiotic resistance is accomplished, we used high-resolution cryo-electron microscopy (cryo-EM) to visualize the structures of ribosomes from both the wild type and the mutant ribosomes, as well as their complexes with antibiotics. Our work shows how the deletion of 4 amino acids in the uL6 protein, which is located 80 and 75 Å away from the aminoglycoside antibiotics (AGAs) and chloramphenicol binding sites, rewrites the conformational landscape of the ribosome affecting inhibition. The structural analysis shows the detailed association of tobramycin near the DC and exposes new binding sites for tobramycin, revealing an allosteric mechanism explaining the differences between this antibiotic and kanamycin, and the molecular mechanism resulting in resistance to AGA and collateral sensitivity to chloramphenicol.

**Figure 1.**
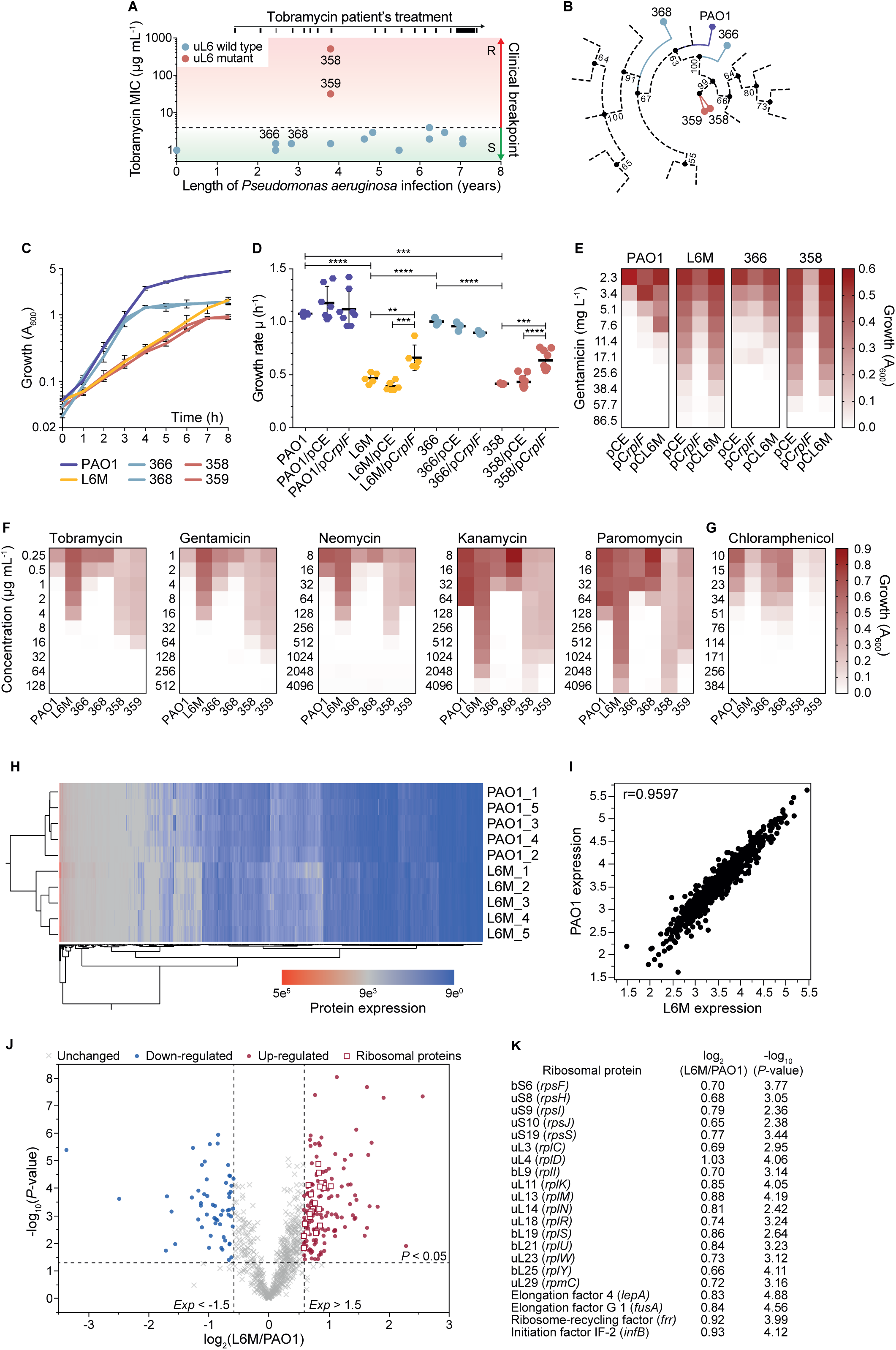
Characterization of *Pseudomonas aeruginosa* strains isolated from CF patient. The different numbers indicate the identities of the clinical isolates. 366 and 368 are the ancestors without the ribosomal mutation while 358 and 359 are the *rplF* ribosomal gene mutants. PAO1 is the wild type of *P. aeruginosa*. L6M is the *rplF* ribosomal gene mutant of PAO1. **A)** Appearance of tobramycin-resistant isolates after repeated tobramycin use. The EUCAST clinical breakpoint for tobramycin is indicated in the graph. **B)** Maximum parsimony phylogenetic tree of the presence-absence of mutations (missense and nonsense SNPs and Indel) in the genomes of the clinical isolates. **C)** Growth profile of the reference strain and the clinical isolates in LB medium. **D-E**) Complementation assay of *rplf* mutation. The wild type and the mutant copy of the *rplf* gene were expressed in the reference and clinical strains of *P. aeruginosa* and the growth rate **D)** and the maximal growth after 48h in the presence of gentamicin **E)** were analyzed. Minimum inhibitory concentrations (MICs) for **F)** aminoglycosides and **G)** chloramphenicol. **H**) Proteome composition of *P. aeruginos*a PAO1 and L6M mutant strains. Hierarchical cluster analysis of the identified proteins. Proteins were clustered according to Euclidean distances using the Ward clustering algorithm^65^. Each biological replicate is represented in the graph. **I**) Correlation analysis of the mean of the expressed proteins in the PAO1 wild-type strain relative to the L6M mutant strain. The Pearson correlation coefficient r is indicated in the plot. **J**) Volcano plots of the proteins showing increased or decreased abundance (*P*-value < 0.05, fold-change >1.5) in the L6M strain relative to the PAO1 wild-type strain. The red and white square represents the ribosomal proteins which expression is increased in the L6M strain relative to PAO1. **K**) List of ribosomal proteins which expression is increased in the L6M strain relative to PAO1. The differentially expressed proteins are listed in Data S1.

## RESULTS

### Identification of a ribosomal mutation causing tobramycin resistance

During eight years of persistent lung infection, clinical strains of *P. aeruginosa* were collected from a CF patient attending the Copenhagen Cystic Fibrosis Center at Rigshospitalet, Copenhagen, Denmark. Antibiotic susceptibility analyses of this strain collection identified two clinical isolates, named 359 and 358, with 8- (359) and 128-fold (358) higher minimum inhibitory concentration (MIC) than the EUCAST clinical breakpoint for the aminoglycoside tobramycin (Fig. 1A). These isolates appeared after intensive treatment with tobramycin and disappeared soon after its interruption (Fig. 1A). Single nucleotide polymorphisms (SNPs), as well as deletions and insertions (indels) relative to the reference lab strain PAO1, were previously identified in these isolates ^22^. Maximum parsimony analysis of the mutations (SNPs and Indels) accumulated in the genomes of the isolates showed a phylogenetic distance of strains 358 and 359 from the other isolates and PAO1 (Fig. 1B). Comparative genomics searching for mutations likely to be involved in aminoglycoside resistance and present only in the resistant strains, revealed an *in-frame* 12-nucleotide deletion (GCTTTGTAACCA) in the *rplF* ribosomal gene of both isolates. This mutation results in a “GYKA” deletion mutant after residue 92, eliminating a protein loop in the C-terminal domain of the uL6 ribosomal protein. The growth rate of isolates 359 and 358 were on average 2.5 ± 0.1-fold lower than that of the ancestor isolates 366 and 368, as well as that of the reference strain PAO1 (one-way ANOVA followed by Tukey’s *post hoc* test; *P* < 0.0001) (Fig. 1C).

To confirm that the *rplF* mutation is responsible for the increase in tobramycin resistance and the reduced growth rate, and to exclude any epistatic interaction from additional mutations present in the genome of the clinical isolates, we created a recombinant derivative of *P. aeruginosa* PAO1 by allelic replacement. The mutant strain, termed L6M, carries the same 12-nucleotide deletion in the *rplF* gene as isolates 358 and 359. Strain L6M showed the same reduction in growth rate relative to that of PAO1 as observed for isolates 359 and 358 (Fig. 1C). We complemented the *rplF* mutation in strain L6M and isolate 358 by heterologous expression of the wild-type copy of the *rplF* gene under its promoter from a plasmid. The results showed a significant increase in the growth rate of 41.1% and 53.2%, for strain L6M and isolate 358 respectively (one-way ANOVA followed by Tukey’s *post hoc* test; *P* < 0.001) (Fig. 1D), and a considerable decrease in the MIC for gentamicin (3.4- and 5-fold respectively strain L6M and isolate 358) (Fig. 1E), in support of the results obtained for the mutant strains. Moreover, both the clinical isolates 358 and 359 and the recombinant strain L6M show higher MIC values than their respective ancestor strains 366, 368, and PAO1 for the aminoglycoside antibiotics tobramycin, gentamicin, neomycin, kanamycin and paromomycin (Fig. 1F). Interestingly, strains L6M, 358 and 359 showed decreased MIC values relative to PAO1, 366 and 368 for chloramphenicol, suggesting a collateral sensitivity phenotype for chloramphenicol directly associated with the mutation in the *rplF* gene (Fig. 1G). No MIC differences were observed for other tested antibiotics (Fig. S1A).

To further assess the occurrence of uL6 mutations in clinical isolates, we compiled and aligned 3310 uL6 sequences from *P. aeruginosa* strains available on the NCBI database. We identified several mutations in the highly conserved region (amino acids 87 to 105) of the uL6 including SNPs and deletions (Fig. S1B). These mutations were identified from strain collections from different countries and infection scenarios including CF patients, leg ulcers and bloodstream infections. This analysis suggests that the generation of mutations in that region of uL6 could be a common mechanism to overcome aminoglycoside antibiotic treatment.

### Ribosome functionality in the L6M strain

In bacteria, the content of active ribosomes is stringently controlled to match the growth rate^31^. Therefore, we tested whether the proteome composition was influenced by the *rplF* mutation performing whole-cell proteomics on the PAO1 and L6M strains. The analysis showed that the L6M and PAO1 strains maintain a high degree of similarity in their proteome composition, as shown by the hierarchical clustering and correlation analysis (Pearson correlation test *r*=0.96, *P* (two-tailed) < 0.0001) (Fig. 1H-I). However, among the differently expressed proteins (fold-change ± 1.5, *P*-value < 0.05), several components of the 30S and 50S subunits were expressed in higher amounts in the L6M strain (Fig. 1J-K, Data S1), suggesting that the mutant bacteria attempt to counterbalance the lower growth rate due to the uL6 mutation by increasing the expression of 30S and 50S ribosomal components. This finding is striking since a reduced bacterial growth rate is usually accompanied by a reduced expression of ribosomal genes ^31^.

### The ribosomes of PAO1 and L6M strains display different configurations

To understand at the molecular level the growth defects, the AGA resistance as well as the chloramphenicol collateral sensitivity mechanisms, we determined the structures of PAO1 (wild type) and L6M (mutant) strains 70S ribosomes by cryo-EM (hereafter termed R and L respectively). In addition, we also determined their complexes with tobramycin, kanamycin and chloramphenicol (Fig. 2A, Fig. S2-S5, Data S2). As datasets in the absence of antibiotics showed lower heterogeneity, we processed them with the local refinement of the three main domains (50S, 30S-body and 30S-head) to reduce the heterogeneity even more without decreasing the number of particles used in the final refinements (Fig. S2-3, Experimental Methods). We obtained excellent density maps and the global resolutions of the structures reported in this work range between 2.4 Å and 2.8 Å, R and L respectively (Data S2). The maps displayed unambiguous densities for all the structures allowing model building and refinement of atomic models. Focused refinement of the 30S improved its map quality, alongside the 30S contacts to the 50S subunit (Experimental Methods and Data S2).

**Figure 2.**
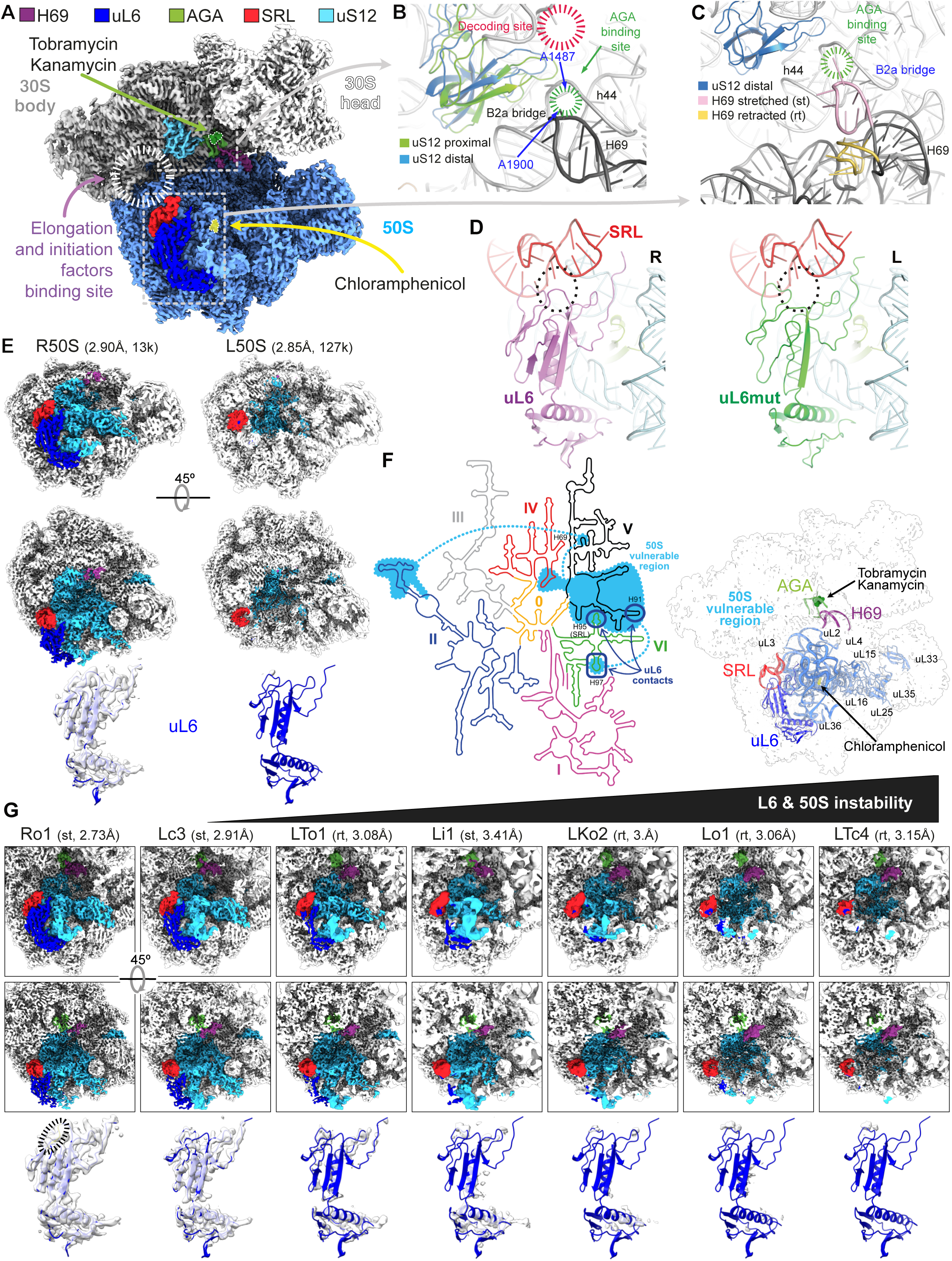
Cryo-EM structural analysis of the R and L ribosomal particles. **A)** Cryo-EM map of *P. aeruginosa* R ribosome without antibiotics (2.4 Å global resolution). This 3D reconstruction of the wild-type ribosome is a composite of the local refinement maps of the 50S, the 30S body, and the 30S head (Fig. S2-3). The positions of the AGA binding site in the h44 helix (green) and the PTC where chloramphenicol (yellow) binds are indicated with dotted white lines as their view is occluded by the ribosome densities. The positions of uL6, uS12, H69, the SRL loop and the region where the initiation and elongation factors associate with the ribosome are also indicated. **B)** Comparison of the uS12 distal (open) and uS12 proximal (closed) configurations observed in the R ribosome in the region where the DC and the AGAs site are located. **C)** Superimposition of the L ribosome shows the differences between the stretched (st) and retracted (rt) H69 helix conformations in the distal configuration. **D)** Comparison of the SRL interaction with uL6 in the R and L ribosomes. The region GYKA-SRL interaction is encircled in a dashed line in the R ribosome, and it is missing in the L. **E)** Cryo-EM maps of the disassembled 50S subunits of L and R reveal that the uL6 protein binding site is distorted. The 50S vulnerable region is depicted in light blue and the uL6, the SRL, and the H69 helix as in A. The density associated with uL6 is indicated in the lower panels, showing that the mutant uL6 is missing in the L50S particles. **F)** Description of the 50S vulnerable region. The left panel shows the RNA secondary structure plot, derived from *E. coli* 23S rRNA, indicating the RNA regions affected by the uL6 instability. The right panel depicts the L 70S particle, showing the links between the uL6 protein docking site in the 50S subunit with the antibiotic binding sites, the H69 RNA helix, the uS12 protein and the 50S vulnerable region. The proteins affected, totally (uL33, uL36, and uL16) or partially, are indicated. **G)** Comparison of the map densities associated with uL6 and the 50S vulnerable region (colors as in E) in different R/L 70S particle reconstructions (both, the conformation of H69 and the global resolution of the maps, are indicated with the name of the class). For each map, the corresponding uL6 protein is represented as an isolated cartoon at the bottom of the figure with its related cryo-EM density (black dashed oval, GYKA). A gradient of stability of those regions of the 50S subunits can be identified from the total stability of the R ribosome to the increased flexibility of the L ribosomes, with the L-50S particle as the extreme case (see Fig. 2E).

The maps show large-scale heterogeneity due to intersubunit rotations between the 30S and the 50S. They display a gradient of conformations, ranging from open to closed states previously described ^7, 32, 33^. The structures are characterized by the distal or proximal positioning of the uS12 protein, and the rest of the 30S excluding the head, with respect to the AGAs binding site in h44 (Fig. 2B, Fig. S2-S5, Data S2, Experimental Methods). uS12 has been proposed to collaborate with the decoding site to optimize codon recognition ^34^, and mutations in two conserved loops made ribosomes substantially resistant to the miscoding effects of paromomycin ^35^. A wide range of conformations between the open/closed states was visualized for R and L ribosomes using a combination of statistical classification and deep neural networks ^36–40^ (Fig. S2). They are similar to previously reported configurations observed in cryo-EM reconstructions ^41–44^ and smFRET studies ^45^. These different conformations are characteristic of the spontaneous intersubunit movements, which permit the oscillation between the two forms, with the equilibrium shifted towards either the open or closed forms, depending on the functional state of the ribosome and the transit of tRNAs during the elongation phase ^7, 41^. To perform a deeper analysis, the different maps were identified as open, intermediate, closed and tightly closed (hereafter named o, i, c and t). In this regard, each reconstruction is named after the purified sample (wild-type or mutant, R or L), the presence of antibiotic (none, Tobramycin, Kanamycin or Chloramphenicol, as ( ), T, K or C), its conformation (o, i, c, t) and an identifying number, i.e., Ro1, rTo1, LKc3, etc. (Experimental Methods).

The structures of the mutant ribosome display a more complex pattern than the wild type, as they combine the open and closed configurations with the “stretched” (st) or “retracted” (r) conformations of the H69 helix in the 23S rRNA^23^ (Fig. 2B-C). The contacts between the 30S and 50S subunits in the B2a bridge between the h44 and H69 RNA helices in the neighborhood of the AGA binding site are different. The H69 helix in the stretched conformation interacts with h44, as in R, while in the retracted conformation the H69 helix is curved towards the large subunit in the distal configuration of uS12. In the R and L stretched configurations, the universally *E. coli* conserved nucleotides A1486 and A1487 in h44 (corresponding to A1492 and A1493 in *P. aeruginosa*), A1900 of H69 and G530 in the DC adopt a similar conformation. However, the sharp kink observed in the retracted H69 helix eliminates the interactions with h44. As a result, the contacts within the B2a bridge region are disrupted, and the A1900 is positioned ∼20 Å away from A1487.

The binding of EF-G to the ribosome in the presence of uncleavable GTP analogs locks the closed form, also termed “rotated” ^33, 45^, while the hydrolysis of GTP and the resulting release of EF-G leads to restoration of the original “non-rotated”, corresponding to the “open” configuration of the ribosomal subunits ^46, 47^. These conformational movements are essential during the elongation phase and for aspects of initiation, fidelity mechanism, termination, and ribosome recycling. Furthermore, the 70S ribosome structure analyses have also shown additional details of configurations in the 30S head domain accompanying the rotation ^48^. Although we were not able to identify translation factors in our maps, they show distinctive distributions of open/close conformational states, and in some cases, tRNAs and mRNA densities can be observed. The deletion of 4 residues in the uL6 of the mutant ribosome could potentially affect the EF-G interaction with the 50S sarcin-ricin loop (SRL) (Fig. 2D). The SRL plays an important role in the stimulation of the GTPase activity of the initiation and elongation factors such as IF2, EF-Tu and EF-G ^49–51^. Likewise, the SRL is also essential for the assembly of the functional core of the 50S subunit ^52^. The lack of these well-conserved 4 residues in the loop produces a shorter connection between the β7 and β8 strands, which destabilizes the secondary structure of the C-terminal antiparallel β-sheet and affects the interaction of uL6 with the SRL nucleotides A2644 to A2648 in the L ribosomes (Fig. 2D).

### The 50S subunit of the mutant ribosomes is misassembled

Interestingly, during the isolation of the mutant ribosomes, a higher number of disassembled subunits were also co-purified when compared with the preparations of the wild type (Experimental Methods). This pattern was confirmed after the analysis of the cryo-EM grids, showing that besides the 70S, the L ribosomes presented many disassembled 50S and 30S particles (Fig. S2B). To investigate possible assembly defects, we also picked the individual 50S and 30S particles and solved the structures of the disassembled subunits (Fig. S2-S5, Data S2). While the 30S subunits did not present any major difference, a comparison of the structures of the 50S subunits of R and L ribosomes, revealed that several proteins, including uL6, and a large section of the adjacent 23S rRNA cannot be modeled in the mutant 50S (Fig. 2E-F). A specific region of the mutant 50S, which we termed the 50S vulnerable region, showed large flexibility, suggesting that the flexibility associated with the mutation in uL6 induces a defective assembly of the 50S particle ^52^. This region includes the SRL of the 23S rRNA, which is contacting the 4-aminoacids loop deleted by the mutation in uL6 (Fig. 2D-E). The RNA helices H91 and H97 also contact uL6 (Fig. 2F). The modification of all these associations seems to expand the effect of the uL6 mutation through the 50S vulnerable region, which connects the assembly site of uL6 with the Peptidyl Transferase Center (PTC) (Fig. 2F), thus supporting the notion that the increased instability observed in the ribosome is generated by the uL6 mutation and transmitted through the 23S rRNA within the 50S, and to the whole ribosome. In fact, a gradient of stability can be observed across the L ribosome maps, where the absence of density corresponding to uL6 correlates with the larger disturbance in the 50S vulnerable region (Fig. 2G, Data S2). Previous studies have also identified this region as a folding block of the 23S RNA ^53^. These findings highlight the importance of uL6 as an essential scaffolding element.

The retracted conformation of the H69 helix also seems to be associated with the growth in the instability of the 50S vulnerable region and the uL6 itself, hence the disassembly of the mutant ribosomes into defective particles like the L50S can be seen as the ultimate consequence of the uL6 mutation and its damaging influence (Fig. 2G). Interestingly, our proteomic analysis showed that the L6M strain overexpresses different ribosomal factors, such as initiation factor IF2, elongation factors EF-G and EF-4, as well as ribosome recycling factors, which bind in the SRL region (Fig. S1J-K, Data S1). This suggests that the L6M strain synthesizes more of these factors to compensate for the effect of the uL6 loop deletion in the assembly of the 50S subunit and the lower amount of 70S functional particles. Collectively, the data indicate that the deletion in uL6 causes instability of the 70S particle conformational dynamics and additional disassembly.

### Analysis of the conformational heterogeneity

Following the initial assessment, we performed an in-depth characterization of the conformational landscape of the different structures with cryoDRGN ^39, 40^. The resultant continuous distribution of particles was divided into 10 clusters per dataset, resulting in 80 initial different classes. This clustering method provides a more accurate distribution of conformations with a higher consistency between different datasets (Experimental Methods, Fig. S2, Data S2). As a first approach to analyze the conformational heterogeneity in the distribution of particles from the latent spaces computed with cryoDRGN, we assessed the degree of variation between the open and closed states, measuring the distance between A55 in h15 of the 30S and G2649 in the SRL in all the different conformations (Fig. 3A inset, Fig. S6A).

**Figure 3.**
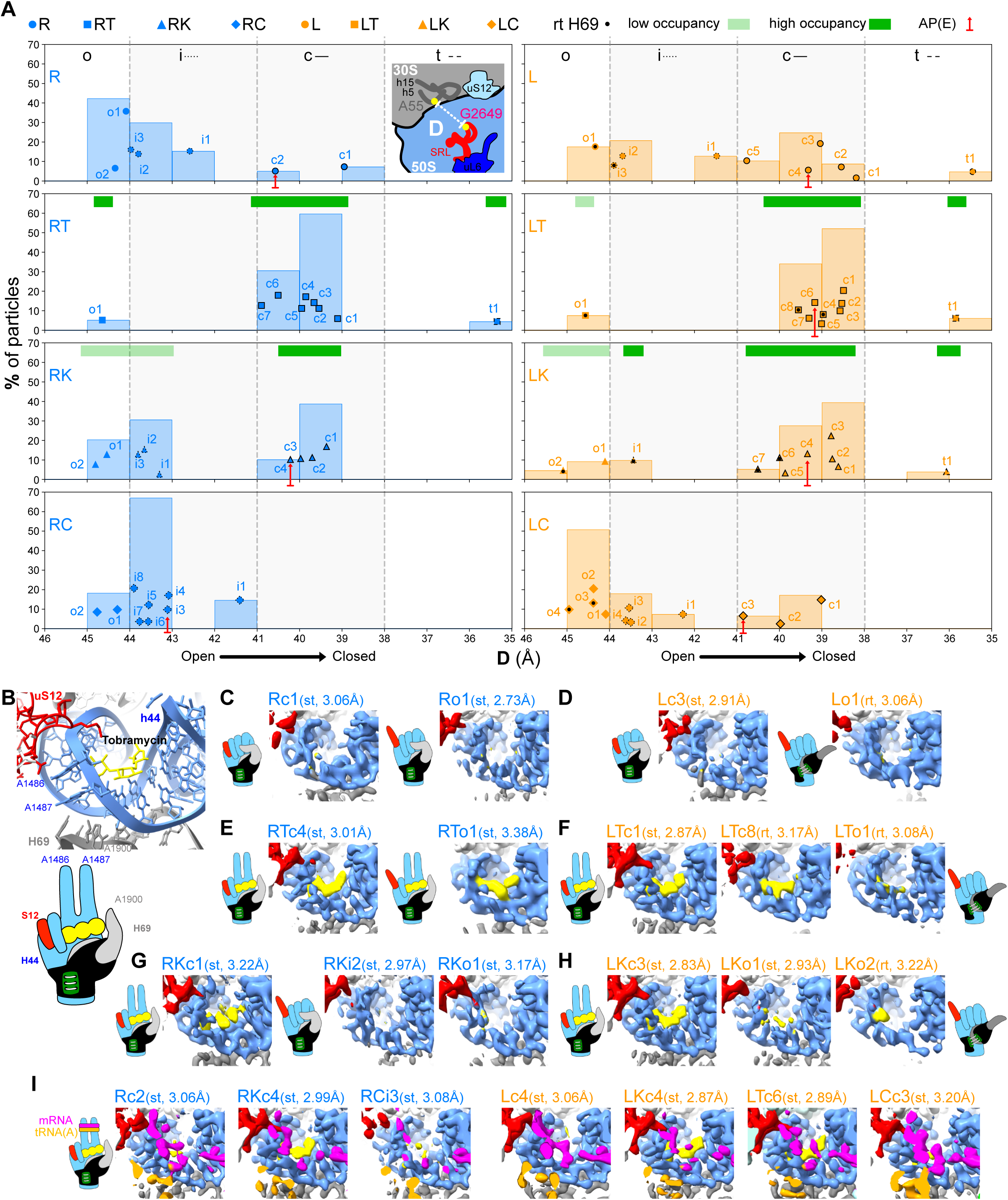
Heterogeneity analysis of the ribosomal particles. **A)** Clustering of the different datasets showing conformational preferences depending on the conditions (R or L purified ribosome, incubated with or without antibiotics). Each class is defined by the distance D and the percentage of particles that were used in its 3D reconstruction (percentage of the total dataset). The sketch in the R histogram defines D (see also Fig. S6A). Binned histograms are included to assist in the interpretation and comparison of the conformational distribution of the particles. To facilitate the identification and description of the classes four distance ranges are defined: o, i, c and t (respectively: open, intermediate, closed, and tight; also indicated in the shape of each datapoint as without line, with a dotted line, with a continuous line, and with a dashed line, respectively). The retracted conformation of helix H69 is shown as a dot at the center of the data points. For the classes incubated with tobramycin or kanamycin horizontal green bars indicate the relative occupancy in the AGA binding site. Red arrows signal the position of those classes with tRNAs in the AP(E) sites (the tRNA at the E site is only partially visible; Fig. S6C). **B)** Detailed view of the AGA binding site. The sketch below of the illustration represents a hand simplifying the different configurations observed in the AGA binding site in the structures of R and L ribosomes. The distal or proximal conformation of uS12, the configuration of nucleotides A1486, A1487, A1900, and the conformation of helix H69 (stretched or retracted) are represented by the fingers. The 50S vulnerable region (Fig. 2F-G) would act as a black wristband which is “tightened” by the uL6 protein (green and white). In R the uL6 protein is always properly assembled in the 50S, thus “tightening up” the hand, while in some cases of the uL6 mutant, its instability can be transmitted and amplified to the 50S vulnerable region and the H69 bridge. **C-H)** The rest of the figure depicts cryo-EM densities of the AGA site for different classes accompanied by a schematic depiction as in 3B. The name of each class is included with an indication of the conformation of its helix H69 and the global resolution of the map. **I)** The panel adds an extra element, two rubber bands keeping the h44 fingers in an extended configuration representing the tRNA (A site) and mRNA interacting near the DC.

The selection of this location in the vicinity of uL6 is related to the rotation movement of the ribosome, as this region is where the EF-Tu translation factor associates with the shoulder of the 30S-body domain, in an open conformation, and its association with the SRL, promotes the rotation towards the closed conformation, leading to GTP-hydrolysis and A-tRNA accommodation ^50, 51, 54^ (Fig. S6B). This distance (D) was used to catalog the 80 different classes obtained from the neural network according to their location in the conformational open-to-closed gradient and analyze the conformational distribution of the different samples (Fig. 3A). This preliminary analysis illustrates the conformational landscape of the intersubunit movements reflected in the different configurations observed by cryo-EM, thus describing the changes in the particle distribution experimented by the ribosome, the effects of the mutation and the influence of the incubation with the antibiotics.

### uL6 mutation and aminoglycoside binding modify ribosome dynamics

The binding of aminoglycosides has previously been reported to impinge on the spontaneous rotation of the ribosome subunits ^45, 55^. The wild-type ribosomes display an asymmetric conformational distribution centered on the open configuration with a decreasing tail into the intermediate and closed conformations. The addition of AGAs produces a shift in the distribution of particles towards the closed conformation. The shift is more homogeneous for tobramycin while the presence of kanamycin favors a more disperse distribution that resembles the wild type in the absence of antibiotics (Fig. 3A). Surprisingly, the mutant in the absence of antibiotics shows already a bimodal distribution, some of the classes behave like the wild type while others shift into more closed conformations. The incubation of the mutant ribosome with AGAs reinforces, even more, this shift, and following the trend shown by the wild type, the effect seems to be stronger for tobramycin (Fig. 3A).

The inspection of the AGAs binding site near the DC reveals that the association of tobramycin and kanamycin is similar in the wild-type and mutant ribosomes (Fig. 3B-I). These antibiotics bind in the h44 helix as previously described for other aminoglycosides ^6, 55, 56^. Both molecules force the flipping out of the A1486 and 1487 nucleotides from the h44 helix to accommodate the antibiotic molecule (Fig. 3B, 3E-I). In the absence of antibiotics, all classes display a disorganized h44 binding site, independently of their conformational state (Fig. 3C-D). In this regard, the cryo-EM densities of both, A1486 and A1487, cannot be unequivocally assigned as their orientation seems to be highly flexible, probably alternating between the flip-out and intercalated states, as there is no perfect RNA sequence match in this site, which is designed for flexibility and insertion of different molecular entities. The occupation of one of the rings of the aminoglycosides into this flexible site mimics the result of the interaction at the DC of a tRNA (A site) and the mRNA, which captures both bases of the helix h44 (Fig. 3I). Consequently, the presence of an AGAs is, in principle, compatible with the formation of the DC, although it seems to affect its function ^6, 55, 56^. Remarkably, the existence in the wild-type and mutant purified ribosomes of particles containing mRNAs and tRNAs (AP(E) classes) allows us to use them as a “molecular ruler” in our distributions of conformations (Fig. S6C, see red arrows in Fig. 3A), as we can use them as internal references to relate the distributions of conformations.

An important observation explaining the different inhibition power between tobramycin and kanamycin is that while the D value of the AP(E) class in the wild type and the mutant ribosomes is almost identical in the presence of kanamycin, the addition of tobramycin eliminates the AP(E) class from the distribution of particles only in the wild type (RT). The AP(E) class is present in the wild type without AGA and the mutant with tobramycin (R and LT, Fig. 3A), suggesting that the presence of tobramycin affects the formation of the AP(E) assemblies more strongly than kanamycin. Furthermore, the L, LT and LK datasets show a consistent shift and apparently, the presence of tRNAs and mRNA in the DC is not affected by the presence of either tobramycin or kanamycin in the mutant (Fig. 3A, 3I, S6C).

### Conformations of the mutant ribosome can escape AGA binding

In the mutant ribosomes, the stretched and retracted configurations of the H69 helix and the instability of the 50S vulnerable region add a layer of complexity to the AGAs binding dynamics (Fig. 2C, 2E-G), leading to different AGA occupancy in the h44 binding site (Fig. 3E-H, Fig. 3A see horizontal green bars). The RT classes display high occupancy of the antibiotic independently of their location in the conformational distribution. In the case of the RK classes, the open and intermediate conformations present low occupancies, as these conformations contain fewer restraints in the AGA binding site (uS12 in distal configuration). This observation highlights the binding and chemical differences between kanamycin and tobramycin. A similar behavior is observed in the LK dataset, where a gradient of occupancy is identified from the closed to the open conformations (Fig. 3H). In the mutant, the addition of tobramycin also produces a class in the open category with low residual occupancy (Fig. 3F). Noteworthy, the retracted conformation of the helix H69 and the state of the vulnerable region could contribute to this situation for the L ribosomes, (Fig. 3A right histograms, rt configuration indicated with a dot). In that sense, while in the L dataset, the rt configuration seems to be restricted to the open-intermediate transition, the addition of AGA introduces also some rt classes in the closed conformation with relatively high occupancy. This observation emphasizes again the differences observed between the two aminoglycosides in wild-type and mutant ribosomes (Fig. 1, Fig. S1).

### The mutant ribosomes are stabilized by chloramphenicol

Chloramphenicol binds directly to the A-site on the 50S ribosomal subunit, occupying the same location as the aminoacyl moiety of an A-site tRNA ^57^. All samples in the presence of chloramphenicol show the drug bound in the PTC (Fig. S6D). Chloramphenicol is bound in a pocket formed by U2491, U2493, C2439, A2438 and G2048. The addition of chloramphenicol to the wild-type seems to concentrate the distribution of conformations into the open-intermediate transition, a smaller change when compared with the effect of AGAs (Fig. 3A, RC and LC). In the case of the mutant, the presence of chloramphenicol seems to produce a distribution of conformations more like the wild type without antibiotics than in any of the mutant datasets. This suggests that the presence of chloramphenicol is reverting the influence of the uL6 mutation in the distribution of conformations (Fig. 3A, LC). This effect is also manifested in the disappearance of classes in the tight conformation (Fig. 3A), which is characteristic of the mutant datasets in the absence and presence of AGA and is also observed in the wild type in the presence of tobramycin. In the AP(E) classes, the presence of the antibiotic induces a considerable shift to more open conformations in the wild-type ribosome (Fig. 3A, 3I), thus displaying the opposite effect of that observed in the AGAs. The LC AP(E) class also experiences a shift in the same direction, but now from the more closed conformation present in the mutant datasets to a D value equivalent to the one for the wild-type AP(E) class (Fig. 3A). Again, the changes in the particle distribution in the presence of chloramphenicol in the 50S compensates the effect of the uL6 mutation. However, it seems that it cannot prevent the disassembly or favor the reassembly of the 50S vulnerable region and the rt configuration of H69 (Fig. 3A, see LC, Fig. S6E).

Collectively, the data indicate that while tobramycin and kanamycin bind and inhibit a representative fraction of the mutant ribosomes, a small population can escape AGA inhibition due to the modification of the conformational changes induced by the mutation. Our results also suggest that stabilizing the chloramphenicol-ribosome promotes enhanced sensitivity and that the effect of chloramphenicol goes beyond blocking the PTC and could be also involved in the conformational dynamics of the 70S particle. However, these local effects around the DC and the PTC do not seem to globally explain the resistance mechanism and its associated collateral sensitivity.

### Disentanglement of the rotational ribosome landscape

Our simple analysis measuring a distance as an indicator between the open and close conformations of the ribosome, revealed the complexity of the conformational detuning of the ribosome dynamics caused by the mutation, showing that the mutant can still be associated with tobramycin. To globally deconvolute and understand the complexity of the conformational landscape, we identified previously defined regions of the ribosome ^58^ behaving as rigid bodies (Fig. 4A). They include three subdomains in the 30S subunit comprising the head and the body, which was subdivided into the B1 and B2 subdomains, and the 50S subunit. Using the B2 domain as a reference, we can study the movements of the other domains in relation to the B2 main elements, the section of the h44 helix where the AGAs bind, the DC and the B2a bridge (Fig. 2B-C). Hence, a comparison of the conformational dynamics was performed by superimposing the different structures in B2 and describing the rotational movements experimented by the B1, the 50S and head domains from a common reference class (Ro1) (Experimental Methods). For each rigid body movement, an associated rotation axis and a rotation value were calculated. Displacement vectors representing the degree of movement of the main chains of the compared classes were also calculated (Fig. 4B).

**Figure 4.**
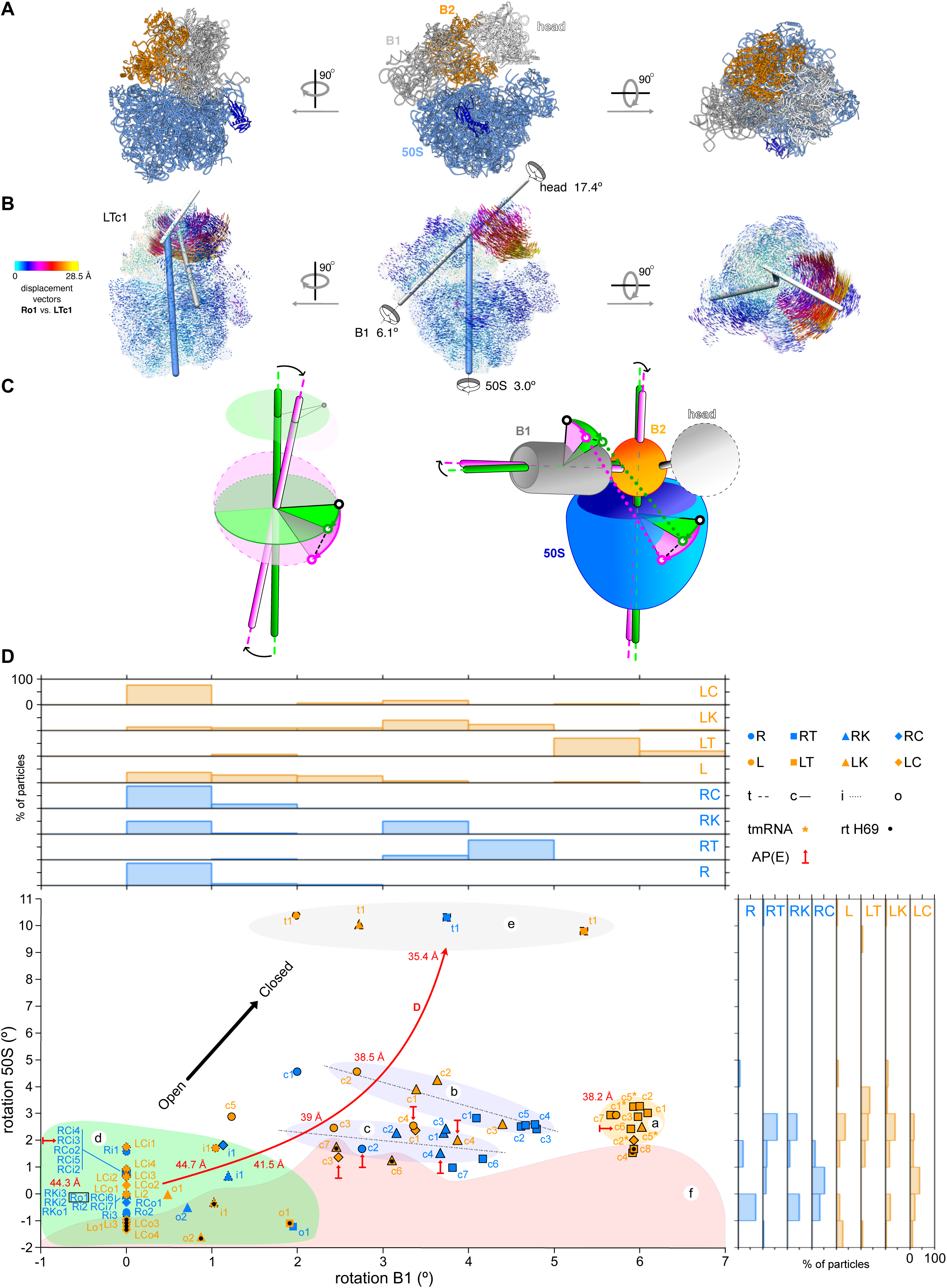
Conformational analysis of the R and L ribosomes. **A)** Definition of the domains Head, B1, B2, and 50S domains used in the rotational analysis. These regions were used to define the axes and rotational displacements for the conformational analyses (Experimental Methods). The image also details the location of the uL6 protein. **B)** Depiction of the domain rotation associated with the transition of the open conformation Ro1 to the closed conformation LTc1. For the calculation of the rotations and displacements the two compared structures were fitted on their 30S-B2 domains, so the movement of the 50S, 30S- B1, and 30S-head can be estimated. For comparison purposes, all rotation axis and values of each class were calculated taking Ro1 as a reference, so these parameters describe the transition from the open conformation of Ro1 to the conformation of a given class. The displacement vectors show the direction and intensity of movement (indicated by the length and color of the arrows) of the domains around the B1, head and 50S axes. **C)** Schematic representation of the effect of changes in the rotational axes (left panel). The position of a given initial point (white circle) is transformed by two different rotations, the axis of one tilted with respect to the other (magenta and green bars, respectively). Rotation planes intersecting in the initial point are depicted as circles with the color of the associated axis. The final coordinates of the point (green and magenta arrowhead) are affected both by the orientation of the rotation axis and the angular magnitude of the applied rotation, depicted as circular sectors in the color of the corresponding axis. The difference between these two positions is represented by a black dashed line. The magnitude of the rotation is also affected by the distance of the initial point to the axis, so the change is amplified as the radius increases (black solid line). The diagram represents a simple case in which the initial point is at the same height as the crossover of the two axes but can be extended to more complicated situations, such as for points at different heights (grey circle and dim upper planes of rotation) or axes with associated translations. The ribosome sketch on the right panel illustrates the effect of two pairs of different rotations, applied to the 50S and B1 domains, associated with two ribosomal conformations (green and magenta). The distance of two initial points after the conformational change (dotted lines) depends on the distance to the rotation axis, the orientation of those axes and the angular magnitude of the rotation. **D)** Plot of the 50S and B1 rotations associated with the transition from the open conformation of Ro1 (reference, zero rotation) to each class. Histogram bins are used to represent the distribution of rotations along the different classes and could be compared with Fig. 3A. Four regions are defined as characteristic bundles of conformational states (see Results). Additional large arrows indicated the direction of the “open to closed” conformational change and approximate values of D (black and red, respectively).

In this scenario, we can consider that the value D is the result of the combination of both the orientation of the rotation axis and the rotation value, with the particularity that the changes will be amplified for regions further away from the rotation axis (Fig. 4C, Experimental Methods). Beyond the detailed characterization of the rotational movements, with variable values and different rotational axes, its interpretation is simplified when plotting just the values of the rotation of B1 versus the values of the rotation of 50S (Fig. 4D), allowing the correlation of different datasets and classes gathered in different clusters that rationalize the original latent spaces from cryoDRGN. This approach deconvolutes the local D values as the result of two concurrent independent rotations describing the conformational dynamics of the ribosome, thus expanding the local structural characterization to a global interpretation (Fig. 3A, 4D).

Almost all mutant structures with tobramycin (LT) locate at the cluster (a), but only two outliers (LTo1, LTt1) (Fig. 4D). Despite having similar D values to some of the LK classes (Fig. 3A), these LT members experience a larger rotation of the B1 domain but conserving the rotation range of the 50S. This can be considered a unique conformational feature of the mutant in the presence of tobramycin, as even the RT classes undergo smaller B1 rotations for similar 50S rotations. Cluster (b) contains some members of RT and LK that share a specific ratio for 50S-rotation/B1-rotation (Fig. 4D, dashed line), where LK can achieve more closed conformations because of larger rotations of the 50S, despite those RT classes experiencing larger B1-rotations. Maintaining this similar ratio implies that as the 50S rotations decrease, the B1 rotations increase, so there is compensation to a certain extent inside the cluster. A similar situation happens with cluster (c). Finally, cluster (d) groups the rest of the open and intermediate classes. Altogether, we can consider the transitions towards the closed conformations as successive jumps from one cluster to the next one. Thus, R classes belong mainly to cluster (d) although more closed classes are located at clusters (c) and (b), which correlate with smaller D values (Fig. 3A and 4D).

The addition of kanamycin promotes a shift of the rotational landscape to cluster (c) and in the presence of tobramycin, almost all classes transit to (c) and (b). In the absence of antibiotics, some classes of the mutant are already shifted towards the closed conformation, thereby the addition of kanamycin shifts them further into cluster (b) and tobramycin induces the jump into the exclusive (a) cluster. As an extreme case, cluster (e) is characterized by larger rotations of the 50S that produce tighter ribosomes (Fig. 3A, 4D, Fig. S7C). Cluster (f) loosely defines an area that contains L classes with the retracted H69 configuration, which displays a larger instability of uL6 and the 50S vulnerable region (Fig. 2G, 4D). In this cluster we also find R classes, which do not show the retracted H69, thus highlighting the fragility of the L ribosomes that cannot hold stably certain conformations.

Despite the heterogeneity and the difficulty when comparing different classes, we can use the ribosomes with densities corresponding with AP(E) tRNAs as a reference to observe the effect of the uL6 mutations and the presence of antibiotics (Fig. 5A-B). When comparing Lc4 to Rc2, the mutant ribosome displays a deviation in the rotation of the 50S, along with a deviation in the rotation of the B1 (Fig. 5A, 4D), producing a different D value (Fig. 3A). The rotation axes of Rc2 and Lc4 cross close to the AGA binding site on helix h44 (Fig. 5A-B). This minimizes the changes in that essential region and the neighboring DC, but the differences are amplified at a greater distance (Fig. 3A, 4C). In addition, as certain components connect different regions of the ribosome (translation factors and tRNAs), whose relative location could be affected by these variations in the rotations, then the fine-tuning of the translation mechanism can be affected. The application of this analysis to the AP(E) classes incubated with antibiotics clarifies the characteristics of the mutant ribosome (Fig. 5C- D). Kanamycin and chloramphenicol produce opposite swivels of the rotation axes when compared to the wildtype class without drugs (Fig. 5C). The inspection of the mutant classes (Fig. 5D) shows that the effect of the antibiotics is added to the own swivel of the mutant (Fig. 5A), thus producing the largest deviation for tobramycin and a compensation for chloramphenicol that makes LCc3 like Rc2. All these changes correlate with the shifts observed previously in the distribution of classes (Fig. 3A, 4D).

**Figure 5.**
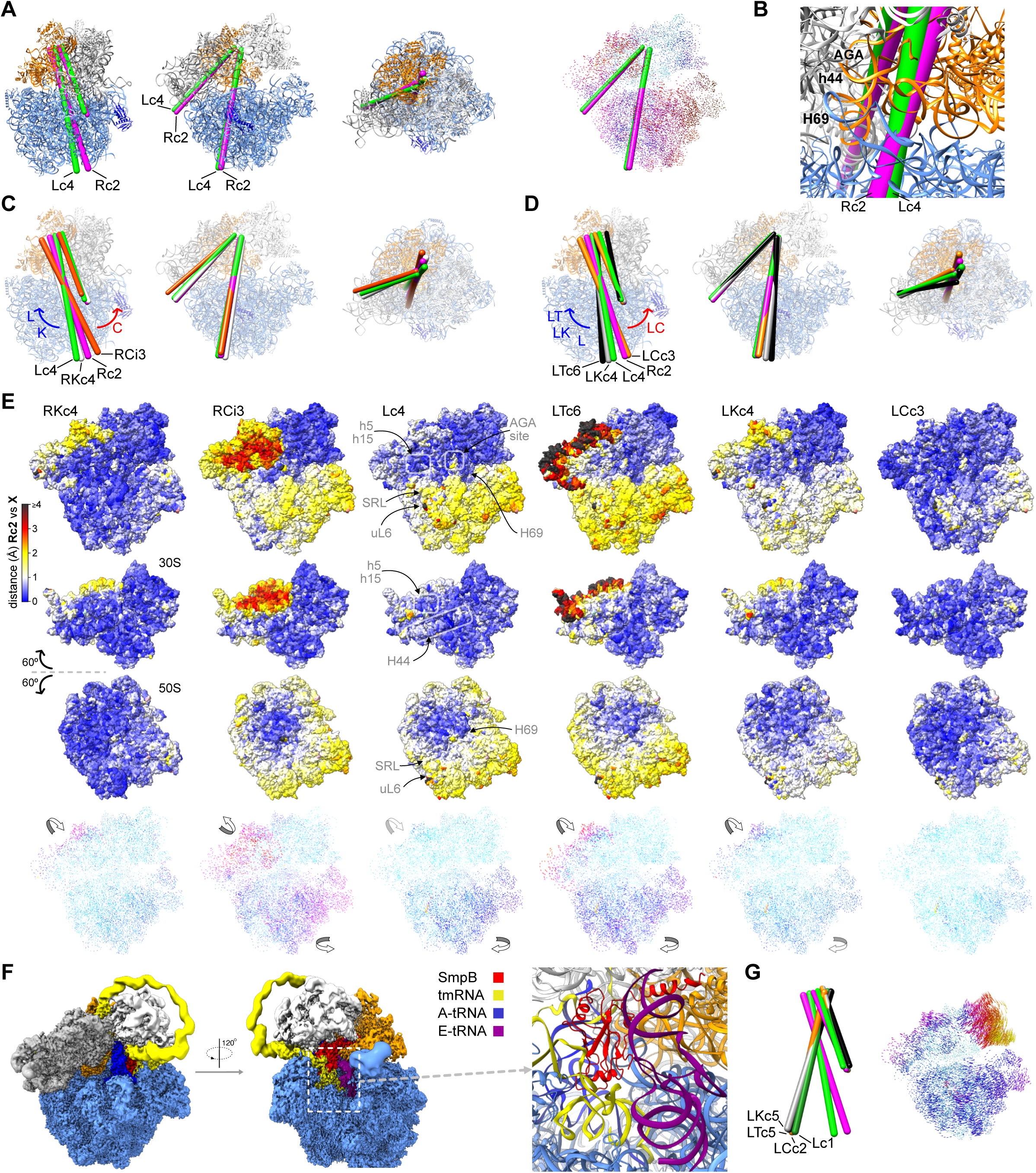
Analysis of the rotation axes associated with the different conformations of R and L ribosomes. **A)** Detailed location of the rotation axes of the Rc2 and Lc4 classes with respect to the structure of the ribosome (reference class, Ro1; same color scheme and views as in Fig. 4A). The inset shows the displacement vectors of Rc2. **B)** Zoom of the first view, rotated 180°, so the closeness of the rotation axes to the helix h44 can be appreciated. **C-D)** Comparison of the rotation axes of R and L classes containing AP(E) tRNAs. Rotations axes of Rc2 and Lc4 are shown again as a reference for comparison of the effects of the mutation and the antibiotics (red and blue arrows). **E)** Analysis of the effects of the previous changing rotational axes on the accessible surface of the ribosomes and their intersubunit interface. Surface representation of the AP(E) models colored according to the distance between equivalent residues (calculated from the Cα and P coordinates, on models fitted by the B2 domain) of Rc2 and the indicated classes. Each model is represented by a view of the 70S particle and 60 [rotations of the 30S and 50S to observe the intersubunit interface. Reference structural elements are indicated in the Lc4 model. **F)** Depiction of the tmRNA-SmpB complex present in the L sample, with a detailed cartoon representation of its components (central panel). **G)** View of the four classes’ rotation axes and associated displacement vectors.

The effect of this rotational behavior affects both the accessible surface and the intersubunit interface of the different classes (Fig. 5C-E). This is visualized when we calculate the deviations between equivalent residues of the classes compared to Rc2 (Fig. 5E). As B2 is used as the reference only minor deviations in this domain are observed, while the full extension of the rotational changes is shown for the B1 and 50S domains. The deviations have a clear radial component, which is better observed in the 50S (Fig. 5E). Changes associated with the uL6 mutation modify the relative position of the 50S but without altering the central intersubunit interface (Fig. 5E see Lc4). In comparison with the 50S, the changes at the B1 domain are smaller but can potentially compensate for the effects on the 50S, thus maintaining a functional ribosome. The binding of kanamycin to H44 induces larger changes at the B1 domain (Fig. 5E, see RKc4 and LKc4). On the other hand, the binding of tobramycin produces the largest changes in both the B1 and 50S domains (Fig. 4D, Fig. 5E see LTc6). In the case of the mutant, these variations compensate for the inhibiting effect of tobramycin in the wild type. As we could not observe an AP(E) class for the wild-type ribosome in the presence of tobramycin, we hypothesize that the binding of this AGA promotes changes incompatible with the assembly of tRNAs and mRNAs. Therefore, the alteration in the 50S displacement introduced by the mutation in L6 would revert this situation allowing translation (Fig. 5C-E. S7E).

Finally, the binding of Chloramphenicol induces relatively large changes in both the 50S and the B1 domains, but in the opposite direction to the ones produced by tobramycin and kanamycin (Fig. 4D, Fig. 5C-E see RCi3 and LCc3). Therefore, the variations introduced by the mutation on the rotation of the 50S and B1 domains are almost completely counterbalanced, making the LCc3 class almost identical to Rc2 (Fig. 5E).

### The uL6 ribosome display classes in complex with tmRNA and smpB

SmpB and tmRNA are involved in ribosome rescue improving the synthesis of full-length proteins and recycling of stalled translation complexes by promoting degradation of truncated polypeptides and turnover of damaged mRNA ^59, 60^. A closer inspection reveals that cluster (a) is not only populated by members of the LT dataset but there are also classes coming from the other three L datasets. The initial purified mutant ribosome, as happened with the AP(E) complex, was also containing a distinctive class of tmRNA-SmpB ribosomal complex after the addition of the antibiotics (asterisks in Fig. 4D, Fig. 5F). These four classes represent a physiological step in the rescue mechanism, with a distinctive feature, together with the tmRNA molecule and the SmpB protein. They contain clear densities for a tRNA in the A-site and low density for a tRNA past the E-site and in the solvent side of the ribosome (Fig. 5F). This implies a close interaction between the A-tRNA and the tmRNA, which is providing the resume codon, and the tail of SmpB occupies the E-site. The presence of all these components promotes a large rotation of the head, accompanied by a rotation of the 50S that is mostly similar for all the classes and swiveled with respect to the previously detailed Lc4 rotation axis (Fig. 5G). As previously mentioned, it is important to emphasize that the LT classes use similar rotations of the B1 and 50S than these tmRNA classes, but with different orientations of the rotation axis. Therefore, the data suggest that the mutant in the presence of tobramycin is reconditioning the movements of the rescue mechanism but with different components in the 50S-30S interface.

### Tobramycin binding sites

The detailed examination of the maps of the mutant and wild-type ribosomes incubated with tobramycin revealed four additional binding sites to the previously described AGA binding site in h44 (Fig. 3B-I, labeled as site 1 in Fig. 6A, S8A). The occupancy of all these novel binding sites can be correlated with the observed conformational behavior of R and L classes (Fig. 3A, 4D). Site 2 is located at the 50S subunit, in the transition of H62 to H63 and close to uL19 and it is close to the intersubunit bridge B6 where helix H62 of the 50S contacts h44 of the 30S (Fig. 6A-B). Only tobramycin can bind to this site, both in L and R classes, as no comparable density is detected in the presence of kanamycin (Fig. 6C). Site 3 lays inside helix h44 at the B6 bridge, close to the uL14 protein (Fig. 6A-B). This site only shows significant densities in L classes in the presence of tobramycin (Fig. 6C). Marginal densities, indicating very low occupancy, can be identified in some RT and LK classes with the degree of rotation of B1 being the main factor favoring binding (Fig. 6, Fig. S8B). Site 4 is positioned between helix h5 and h12, close to uS12 (Fig. 6A-B). It shows the same occupancy pattern as site 2, exclusive of tobramycin but for both R and L classes (Fig. 6C). Site 5 is located at helix h7, close to uS16 (Fig. 6A-B) and behaves as sites 2 and 4 (Fig. 6C). As expected, in the presence of only chloramphenicol all these sites show no significant densities (Fig. S8C). Although other secondary binding sites have been documented for other aminoglycosides at the basis of the H69 helix, i.e. neomycin ^55^ and paromomycin ^10^, we have not been able to detect tobramycin or kanamycin in that location at the concentrations used in this study (Experimental Methods).

**Figure 6.**
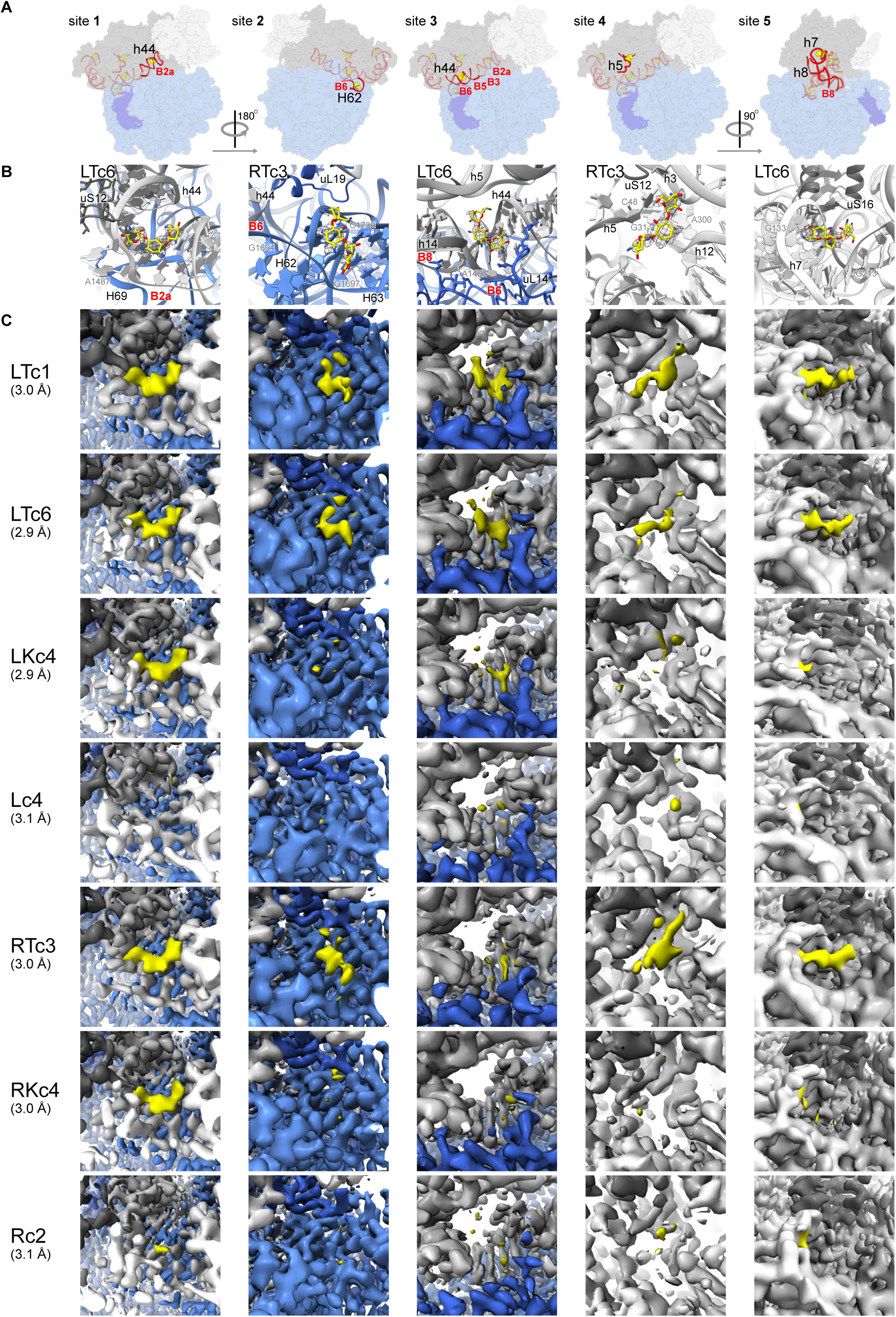
Tobramycin binding sites in the wild type and mutant ribosomes. The figure is organized in columns comparing the five tobramycin binding sites between the different datasets. **A)** Sketches of the location of the five binding sites of Tobramycin in the R/L ribosomes, highlighting the main closer structural elements and related intersubunit bridges (labeled black and red respectively; 50S blue, 30S-body grey and 30S-head white). **B)** Detailed view of the five binding sites, including the antibiotic density shown as a mesh with the corresponding map indicated at the top-left corner. **C)** Comparison of the Cryo-EM densities of LTc1 and LTc6 with the other five maps, showing the differences in the occupancy of the binding sites. The color code is defined as in B, with densities corresponding to tobramycin or kanamycin in yellow.

Collectively, these observations explain the different inhibitory actions of tobramycin and kanamycin. The finding of several binding sites for the clinical antibiotic provides chemical support to the singular conformational landscape of the wild-type and mutant ribosomes in the presence of tobramycin and reveals how the mutant turns the antibiotic into an allosteric activator.

## DISCUSSION

We analyzed the phenotype and the molecular basis of a tobramycin-resistant mutant strain of *P. aeruginosa* derived from a CF patient (Fig. 1). Different mutations in the same region of the *rplF* gene were identified in clinical collections spanning different countries and infection scenarios (Fig. S1B), indicating that the selective pressure imposed by aminoglycoside treatment might select and enrich for uL6 mutations. The mutation produces a minor change in the ribosome structure, which generates a non-linear impact on the complex conformational choreography of the ribosome particle. Chemical modifications in the *Saccharomyces cerevisiae* ribosome have been suggested as the cause of different conformations reflective of abnormal intersubunit movements, which lead to deficiencies in translation ^61^. Our study shows that the deletion of the GYKA loop in uL6 also alters the conformational landscape of the ribosome, leading to a destabilization of the C-terminal half of the protein, which displays larger flexibility in the mutant (Fig. 7A, 2D-G). This increase in flexibility influences the assembly of the 50S subunit, where large regions of the 23S rRNA and many proteins are not visualized (Fig. 2E-G). The data suggest that a deficient docking of the uL6 protein due to the lack of stabilization of the SRL and the adjacent RNA contacts is the origin of this problem (Fig. 7A, 2F). The assembly of the uL6 protein seems to be a prerequisite to stabilizing this region, thereby assuring the proper assembly of other factors in the 50S ^52, 53^. Hence, a defective or inefficient uL6 assembly could lead to an unstable particle through the 50S vulnerable region (Fig. 2E-G, 7A). These unstable particles would lose the intersubunit B2a (h44-H69) bridge, promoting ribosome disassembly (Fig. 2E, 2G). On the other hand, an assembled but unstable uL6 mutant would unbalance the vulnerable region and the 50S-30S bridges to induce a detuning of the 70S translation mechanism affecting the conformational behavior without compromising cell viability (Fig. 7B, 2G, 3A, 4D). A comparison of the particle distributions between different conformations of the wild type and the mutant ribosomes indicates that the uL6 deletion decreases their preference to populate the open ribosome conformation, shifting into more closed configurations (Fig. 7B, 3A, 4D).

**Figure 7.**
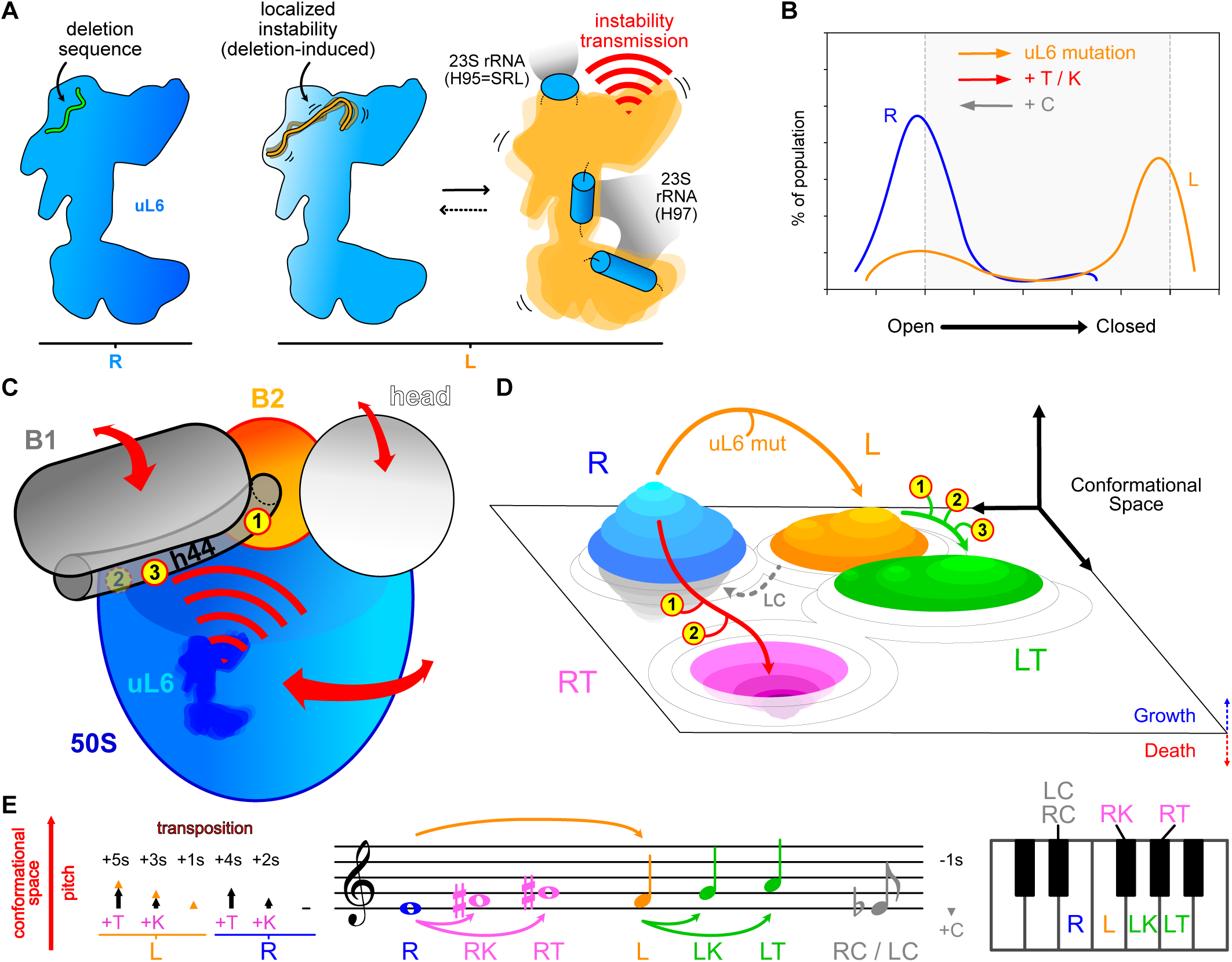
Mechanism of aminoglycoside resistance and chloramphenicol collateral sensibility due to the deletion in the uL6 protein. **A)** Structural instability associated with the deletion of the GYKA loop in uL6. **B)** The presence of antibiotics and the uL6 mutation disturb the conformational landscape of the ribosomes. While the mutation and the AGAs modify the distribution of conformations in the same way, chloramphenicol operates in the opposite direction. **C)** The uL6 mutation affects the internal configuration of ribosomes modifying their conformational behavior in a similar way as tobramycin affects it by binding to different sites (only three out of five sites are shown for clarity). **D)** Exclusive binding of tobramycin to site 3 in the mutant produces a conformational drift that allows overcoming the deleterious effect of the antibiotic. **E)** The Well-Tempered Ribosome. The sketch depicts the analogy between the additive behavior of the conformational changes described in this work (population shifts and domain rotations) and the harmonization and transposition of notes in western classical music. While kanamycin produces a transposition up of two semitones (represented as +2s) and tobramycin of four semitones, the uL6 mutation generates a pitch shift of one semitone. On the other hand, chloramphenicol induces a transposition down of one semitone. The additive nature of these changes makes that while the notes associated with RK and RL are sharp, black keys in a piano, the R, L, LK and LT notes lay in white keys. The latter group always sounds in harmony, as they belong to the C major scale, but the former notes are dissonant to them. Following this analogy, the bacteria can improvise with the ribosome as long as it remains in key, and the melody harmonizes with the rest of the ribosomal mechanisms and functions. In this context, the emergence of the uL6 mutation can be seen as a modulation of the key, from C major to its relative minor, A minor (which provides the mutant with its own feeling).

### The L6 mutant rewrites the ribosome conformational landscape

AGAs bind preferentially in the h44 helix of the 30S subunit near the DC and are supposed to inhibit translation by hindering translocation through stabilization of the 70S, locking intersubunit rotation ^9, 62^. The h44 helix plays a fundamental role in both the sensitivity of the wild-type ribosome to tobramycin and kanamycin and the resistance exhibited by the uL6 mutant (Fig. 7C). In the open conformation, the classical AGA binding site (site 1), is not as confined as in the other structures, and the increased flexibility helps in the adoption of the retracted configuration of H69 (Fig. 2B-C), and the lower occupancy of the h44 binding site for kanamycin and tobramycin (Fig. 3C-D). The incubation with aminoglycosides induces a shift in the conformational distribution of the mutant particles, taking the retracted H69 configuration in some of the closed conformations (Fig. 7B, 3A, Fig. 4D). This configuration of H69 is absent either in mutant ribosomes without aminoglycosides or in the mutant ribosome with chloramphenicol. Collectively, the data indicate that these new classes are produced by the binding of the aminoglycoside to preexisting open-retracted conformations that are shifted to more closed configurations (Fig. 3A, Fig. 4D). Remarkably, we found three new binding sites in the wild-type ribosome for tobramycin besides the AGA site in h44 near the DC (Fig. 6, 7C). However, in the case of kanamycin, which displays a much lower MIC (Fig. 1F), the antibiotic is only present in site 1, both for the wild-type and mutant ribosomes.

The presence of several binding sites of tobramycin in the ribosome has been observed biochemically ^63^. Yet, to our knowledge, no detailed structural study of the inhibition of tobramycin in the bacterial ribosome has been previously performed and the only structural information available was from a crystal structure of tobramycin bound to an oligonucleotide mimicking the h44 helix ^64^.

Furthermore, our analysis reveals that the reshuffling of the conformational landscape of the ribosome by the deletion in uL6 favors the binding of tobramycin in a 4^th^ new additional site (site 3) located at h44. This novel site is exclusive for tobramycin and not observed in the wild type. The new binding sites, especially the one exclusive of the mutant ribosome, explain why the conformational shift towards closed configurations is larger in the case of tobramycin compared to kanamycin (Fig. 7C-D, 3C, Fig. 4C), as the larger extent of these movements can affect the interaction of translation factors and tRNAs with the ribosome. In this way, tobramycin, and perhaps kanamycin to a lower extent, would interfere with the codon-anticodon interaction by bypassing the initial more open conformations, where those regions in the neighborhood of the DC and uS12 remain in a conformation capable to accept a tRNA at position A. However, regions located further away experience larger changes having a larger functional impact. Therefore, to circumvent the inhibitory action of tobramycin, the uL6 deletion would alter the stability of the 50S subunit, generating the additional binding of tobramycin in the novel site3 at the ribosome (Fig. 6, 7C and 7D). This binding would induce a conformational drift, exclusive of the mutant ribosome in the presence of tobramycin (LT), counteracting the harmful effects of the other sites also present in the wild type with the antibiotic (RT). Thus, the mutation in uL6 would convert the antibiotic into an allosteric effector (Fig. 7D). This notion is supported by the presence of tRNAs and mRNAs in the ribosome, which is observed in all cases but on the wild type with tobramycin (Fig. 3A, 4D, S6C). This could explain why its inhibitory action is stronger than kanamycin, reinforcing the idea that the additional binding sites of tobramycin collaborate to alter the conformational dynamics. The L6 mutation rewrites the scenario by allowing the assembly of tRNAs and mRNA with the ribosome favored by the generation of a new binding site for tobramycin (site 3), which is exclusively present on the mutant ribosome (Fig. 6).

### Chloramphenicol collateral sensitivity

The increased chloramphenicol sensitivity of the mutant is also explained by the conformational shifts observed in the mutant ribosome (Fig. 7B, 3A, 4D). In the presence of chloramphenicol, both R and L shift their conformational distribution towards more open configurations, in such a way that in the case of the mutant ribosomes, their distribution looks more like the wild type than that of the mutant ribosomes without antibiotics (Fig. 3A, 4D). As chloramphenicol binds to the PTC in the 50S vulnerable region, it would compensate for the disrupting effects of the uL6 mutation, stabilizing the 50S and by extension the 70S particle, making energetically favorable its binding to the PTC and thereby the increased sensibility to chloramphenicol (Fig. 1G, Fig. S6D). Collectively, the changes that promote resistance to aminoglycosides increase on the other hand the sensitivity to chloramphenicol by stabilizing the mutant ribosomes, as the fraction of particles with the flexible H69 retracted conformation is reduced in the presence of this antibiotic (Fig. 2B-C).

### The side effects of the uL6 mutant

Although the mutant ribosomes can overcome the toxic effects of tobramycin, the growth rate is compromised by the intrinsic instability of the mutant uL6 and the potential misfunction and disassembly of the ribosomal particles. This is supported by the presence of tmRNA-SmpB exclusively in the mutant ribosome samples (Fig. 4D, 5F-G). Although it is difficult to determine whether the presence of the tmRNA complex in the L sample is related to a higher occurrence of stalled ribosomes, a preference of the L ribosome for this intermediate conformation of the rescue mechanism, or if there is some kind of problem that block the resolution of tmRNA complex. Our data indicate that to compensate for the lack of fully functional ribosomes and the lack of assembly of the 50S subunits, the cells increase the expression rates of ribosome and translation factors (Fig. 1H-K).

### The well-tempered ribosome

All the described conformational changes seem to be operating on the very same underlying common mechanism, producing interference when combined, i.e., in the case of the mutant in the presence of tobramycin (Fig. 7C-D). This addition effect can be illustrated with an analogy based on musical composition (Fig. 7E), emphasizing that in the context of a larger set of functional ribosomal conformations and compositions, some configurations are allowed (they are in tune, like L, LK and LT) but others are deleterious (dissonant, like RK, RT, RC, and LC). In this sense, the additional changes introduced by the mutation correct the effects of AGAs, re-tuning the ribosome. Following our interpretation, when comparing kanamycin with tobramycin, the highest binding promiscuity of the latter seems to be responsible for the enhanced sensitivity in the wild type and its purpose as a natural antibiotic product (extracted from *Streptoalloteichus tenebrarius*). However, when confronted with the limitless ingenuity of bacterial natural selection, that very same feature becomes a potential weakness as it can be exploited to revert its initial purpose (Fig. 7C-D). In principle, a modified version of tobramycin could exclusively prevent the binding to site 3 while conserving the other sites, and thus reverse resistance. Yet, the question is how fast the bacteria would circumvent the action of the new molecule, leading to an endless cat-and-mouse game. This natural selection between sensitivity and resistance, would be achieved by chance just by bacterial survival in the lung. Another possibility is that the mutant is just reverting to or repurposing an archaic solution deeply encoded in the ribosome, as a result of countless generations through precursory ancient mechanisms or previous encounters with toxin-producing microorganisms. This information may be used for the development of new antibiotics or the combination of the aminoglycoside treatment with other compounds that target the effects of the mutations to restore the antimicrobial effect.

## SUPPLEMENTAL INFORMATION

Supplemental Data Information includes supplementary figures (S1-S8) and DATA S1 and S2, which consist of two Excel files.

## ACKNOWLEDGEMENTS

We thank Rasmus Marvig for searching for mutations in ribosomal protein genes in a large collection of genome-sequenced patient isolates of *P. aeruginosa.* The Novo Nordisk Foundation Center for Protein Research and The Novo Nordisk Foundation Center for Biosustainability are supported financially by the Novo Nordisk Foundation (Grant NNF14CC0001 and NNF10CC1016517). This work was also supported by the cryo-EM (Grant NNF0024386), cryoNET (Grant NNF17SA0030214), Distinguished Investigator (NNF18OC0055061) grants and ERC_AdG_101096548 INTETOOLS to GM. The Danish Council for Independent Research (Grant Rammebevilling DFF-4181-00115 and DFF-9040- 00106A) and the Cystic Fibrosis Foundation (Grant MOLIN18G0) to SM. AJF was a recipient of a H.C. Ørsted COFUND Postdoc Fellowship from the Technical University of Denmark. We thank the Danish cryo-EM National Facility in CFIM at the University of Copenhagen for support during data collection and Irina Poznyakova for mass spec analysis. Data processing has been performed in the Computerome, the Danish National Computer for Life Sciences. Guillermo Montoya is a member of the Integrative Structural Biology Cluster (ISBUC) at the University of Copenhagen.

## AUTHOR CONTRIBUTIONS

HKJ created the biobank of clinical *P. aeruginosa* isolates from which the original strains harboring the L6 mutation was isolated. SM was the original initiator of the ribosome project. AJF, RLR, REP designed and performed the biochemical and microbiological experiments. REP and PM prepared the cryo-EM samples and PM prepared the grids, collected the cryo-EM images and solved and refined the cryo-EM structures. P.M. and G.M. proceeded with cryo-EM structure analysis. The global results were discussed and evaluated with all authors. G.M. and S.M. coordinated and supervised the project and G.M. wrote the manuscript with input from PM and the rest of the authors.

## DECLARATIONS OF INTEREST

Guillermo Montoya declares that is a member of the SAB and a stockholder of Ensoma. The rest of the authors declare no competing financial interests.

## RESOURCE AVAILABILITY

### LEAD CONTACT

Further information and requests for resources and reagents should be directed to and will be fulfilled by the lead contact, Guillermo Montoya.

### MATERIALS AVAILABILITY

All reagents generated in this study are available upon request to the lead contact with a completed Materials Transfer Agreement.

### DATA AND CODE AVAILABILITY

- The atomic coordinates and cryo-EM maps included in this study have been deposited in the Protein Data Bank and Electron Microscopy Data Bank under the accession codes: **(Given the large number of structures in this study we are in the process of submission. They are available upon request)**
- This paper does not report original code.
- Any additional information required to reanalyze the data reported in this paper is available from the lead contact upon request.

### QUANTIFICATION AND STATISTICAL ANALYSIS

The statistics for the reported cryo-EM structures shown in Table S2 were obtained from Relion 3.1 (overall and local resolution) and Phenix (refinement and validation).

## EXPERIMENTAL METHODS

### Bacterial strains and media

*P. aeruginosa* clinical isolates were previously sampled and identified from sputum samples of patient P76M4 attending the Copenhagen Cystic Fibrosis Center at Rigshospitalet, Copenhagen, Denmark ^22^. The *P. aeruginosa* laboratory strains PAO1 was used as reference. The strain L6M was constructed by replacing the wild-type copy of *rplf* gene of PAO1 with the mutant *rplf* gene from isolate 358. *rplf* gene was amplified by PCR from gDNA of clinical isolate 358 with primers EcoRI-rplF-fwd (TATAGAATTCGTTGATCGCGACATCGAAT) and BamHI-rplF-rev (TATAGGATCCTTCCATTGCTTTCTGGATGG) and cloned into EcoRI-BamHI-pSEVA612S restriction sites ^66, 67^. The resulting vector was transferred by triparental mating to PAO1, using pRK600 as helper ^68^. For the triparental mating, *P. aeruginosa* strain was grown at 42 °C and *E. coli* strains at 30 °C. The first event of recombination was selected by Pseudomonas Isolation Agar plates containing 25 mg L^-1^ of gentamicin. The second event of recombination was induced by Sce-I chromosome cleavage after triparental mating of pSW-I plasmid and induction with 15 mM 3-methylbenzoate ^67^. The presence of the 12-nucleotide deletion was tested by PCR and sequencing. Complementation assays were performed by analyzing the growth profile and the turbidity (A_600_) after 48 hours of growth in the presence of gentamycin of *P. aeruginosa* strains containing the pSEVA332 derivative plasmids, pCE (empty vector), pC*rplF* (harboring the wild-type copy of *rplF* gene) and pCL6M (harboring the mutant *rplf* gene from isolate 358). Complementation strains were created by amplifying, from *P. aeruginosa* PAO1 and clinical isolate 358, the region containing the promoter and *rplf* gene with primers EcoRI_rplF_fwd and HindIII_rplF_rev (TATAAAGCTTATGACCTGGGCGTAAATGTG), cloning it in the pSEVA332 plasmid ^69^ using EcoRI and HindIII restriction sites and by transferring the resulting plasmids to *P. aeruginosa* receptor strains via tripartite conjugation using the pRK2013 helper plasmid ^70^. All PCRs were performed with Phusion Hot Strart II High-fidelity DNA Polymerase (Thermo Fisher). Bacterial growth was recorded by measuring the turbidity at 600 nm of cell cultures in a 250 ml flask at 250 rpm at 37°C in LB medium. For each strain, at least three independent biological replicates were analyzed.

### Antibiotic susceptibility tests

The Minimum Inhibitory Concentrations (MICs) for tobramycin was measured by E-tests (Liofilchem®, Roseto degli Abruzzi, Italy) according to the manufacturer’s guidelines. MICs for aztreonam, carbenicillin, ceftazidime, chloramphenicol, ciprofloxacin, colistin, erythromycin, gentamicin, kanamycin, neomycin, paromomycin, tetracycline, and tobramycin were determined in 96-well microplates in LB medium containing decreasing concentrations (1.5- or 2-fold dilution series) of each antibiotic. Plates were incubated for 48h at 37°C at 150 rpm. For each strain, three independent biological replicates were analyzed.

### Comparative genomics and whole genome sequencing

Genomic data consisting of SNPs and indels are available for the different clinical isolates ^22^. We performed Maximum Parsimony (MP) analysis to highlight differences in the presence and distribution of mutations (missense and nonsense SNPs and indel) between the genomes and to infer the evolutionary trajectory of the bacterial isolates. The bootstrap consensus tree is inferred from 1000 replicates. The analysis involved a matrix of presence-absence of missense and nonsense SNP and indel mutations consisting of a total of 74 mutated genes for each of the 18 isolates and PAO1 reference strain. Evolutionary analyses were conducted in PAUP version 4.0a (build 161). Whole genome sequencing of strain L6M was performed to confirm the 12 nt *rplF* gene deletion and to identify any other mutation on ribosomal RNA or ribosomal proteins present in the genome. The full list of mutations is reported in Supplementary Table S2. L6M strain genomic DNA was extracted using the DNeasy Blood & Tissue kit (Qiagen), the library prepared using the KAPA HyperPlus kit (Roche) and sequenced by NextSeq system (Illumina) with a mean coverage of 100x. The resulting paired-end reads were trimmed with trimmomatic (version 0.38) ^71^ and quality controlled using FastQC (version v0.11.8). SNP and Indel mutations were identified with Breseq pipeline using *P. aeruginosa* PAO1 (RefSeq NC_002516.2) as reference genome ^72^. Sequencing data are available at Sequence Read Archive (SRA) accession number PRJNA530110.

### P. aeruginosa ribosome purification

Precultures of *P. aeruginosa* cells were diluted (1:50 or 1:100 depending on the growth rate) and regrown in LB at 37°C until exponential phase (0.4 - 0.6 A_600_), harvested by centrifugation at 8000 g and 4°C, washed twice in buffer A (50 mM Tris HCl pH 7.6, 10 mM MgCl_2_, 100 mM NH_4_Cl, 0.5 mM EDTA pH 8, 6 mM β-mercaptoethanol) and concentrated 1:100 in buffer A. Cellular lysis was carried out as previously reported ^73^ by adding 1.3 mg mL^-1^ lysozyme, fast freezing and ice-thaw. The lysate was centrifuged at 22000 rpm for 30 min at 2°C and the supernatant was stored at -80°C until further analysis. Ribosome concentration was measured using the O.D. at 260 nm of the preparation.

### Sample Preparation for Proteomic Analysis

*P. aeruginosa* overnight cultures were diluted in M9 minimal medium with trace elements supplemented with 25 mM of succinate and grown at 37°C with 180 rpm until mid-exponential phase (0.4 of optical density at 600nm). Cells were collected by centrifugation, snap-frozen on dry ice and stored at -80°C until further use. Cell preparation and proteomic analysis were performed as previously described ^74^. Briefly, 200 µL of urea (8 M, 75 mM NaCl, 50 mM Tris-HCl, pH 8.2) and two 3-mm zirconium oxide beads (Glen Mills, NJ, USA) were added to the cell pellets. After 15 minutes at 4 °C, cells were disrupted using a Mixer Mill (MM 400 Retsch, Haan, Germany) for 2 min at 25 Hz. After an additional 15 minutes at 4°C, cells were again subjected to 2 min of disruption, after which the samples were centrifuged at 14,000 *g* for 10 min at 4°C. 100 µL of supernatant was collected and diluted with 900 µL of 50 mM ammonium bicarbonate. Samples were then concentrated to 100 µL using a 3 kDa cut-off filter (Amicon Ultra filter, Merk, Germany). For each sample, 100 µg of proteins were incubated at 37°C for 45 min after the addition of 5 µL of 100 mM DTT, followed by a further incubation of 45 min in the dark after the addition of 10 µL of 100 mM iodoacetamide. The samples were digested with 1 µg/sample of trypsin for 8 h, after which 5 µL of 10% TFA was added. Samples were StageTipped using C18 (Empore, 3M, USA) according to a previously described procedure ^75^

### NanoUPLC-MSE Acquisition

For Nanoscale LC analysis of the trypsin-digested samples, a nanoACQUITY system (Waters, USA) equipped with a Symmetry C18 5-µm, 180 µm x 20 µm precolumn and a nanoACQUITY BEH130 C18 1.7-µm, 75 µm x 250 mm analytical reversed-phase column (Waters, USA) was used. For each sample, 1 µg of protein was trapped on the precolumn using mobile phase A, consisting of 0.1% formic acid in water with a flow rate of 8 mL min-1 for 4 min. Mobile phase B consisted of 0.1% formic acid in acetonitrile. A reverse-phase stepped gradient was used to separate peptides: I) from 6% to 14% acetonitrile in water over 28 min, II) from 14% to 25% acetonitrile in water over 40 min, III) from 25% to 38% acetonitrile in water over 15 min, IV) from 38% to 60% acetonitrile in water over 10 min and V) from 60% to 99% acetonitrile in water over 20 min. Between each injection, a 30 min wash method was applied. Both methods used a constant column temperature of 35C and a flow rate of 250 nL min-1. The described gradient data were acquired using a Synapt G2 (Waters, Manchester, UK) Q-ToF instrument operated in positive mode using electrospray ionization with a NanoLock-spray source. Using the internal fluidics system of the mass spectrometer, leucine enkephalin was used as a lock mass. The lock mass channel was sampled every 60 s. For each injection, the mass spectrometer was operated in resolution mode, with continuum spectra being acquired. During acquisition, the mass spectrometer alternated between low- and high-energy modes using a scan time of 0.8 s for each mode over 50–2,000 Da. In the low-energy MS mode, data were collected at a constant collision energy of 4 eV. In the elevated-energy MS mode, the collision energy was increased from 15 to 40 eV.

### Protein Identification and Quantification

Protein identification and quantification were obtained using Progenesis QI for Proteomics version 2.0 and the *P. aeruginosa* UniProt proteome database (ID: UP 208964). Settings for the PLGS search engine were FDR 1%, tryptic peptides with one missed cleavage allowed, and carbamidomethylation of cysteine residues as fixed modification and oxidation of methionine residues as variable modification. For quantification, only unique peptides of the proteins of interest were used, enabling comparisons of protein abundance across the different samples ^76^. Data are presented as the means of 5 independent replicas. The abundance was calculated based on the levels of the proteins in the PAO1 wild type strain relative to the L6M mutant strain using the Anova statistical test. Only proteins with at least a 1.5-fold change relative to the wild-type strains (*p* > 0.05, Anova) and with 3 identified unique peptides were reported as the altered proteome.

### uL6 protein alignment

uL6 protein sequences were obtained from NCBI using the terms L6 AND “Pseudomonas aeruginosa” and further aligned using the Clustal Omega version 1.2.4 algorithm using the software SnapGene 6.2.2.

### Cryo-EM sample preparation and data acquisition

The R and L ribosomal samples were isolated as described above. In addition, for Cryo-EM analysis, those samples were applied onto a CaptoCore 700 column (Merck) for additional purification in buffer B (50 mM Tris HCl pH 7.6, 10 mM MgCl_2_, 100 mM NH_4_Cl, 6 mM β- mercaptoethanol). The column flowthroughs were flash-frozen in liquid nitrogen and stored at -80° C. These samples (R and L) were initially screened on a Tecnai G2 20 TWIN (FEI, Thermo Fisher Scientific) to optimize the appropriate ribosomes working concentration, buffer, grid type and homogeneity. Each sample was diluted to 400 nM in buffer B, in the presence or absence of antibiotics (50 μM final concentration), and then incubated on ice for 20 min. A volume of 3ul of the sample was applied onto glow-discharged (Leica EM ACE200, 30 s at 5 mA) grids (Quantifoil R1.2/1.3 200 mesh Cu grids) and plunge-frozen into liquid ethane, cooled with liquid nitrogen, using a Vitrobot Mark IV (FEI, Thermo Fisher Scientific) with the following settings: 100% humidity, 277 K and blot time of 3 s. Grids for the eight conditions (R, RK, RT, RC, L, LK, LT and LC) were prepared in the same day under common conditions and using the same antibiotic stocks and dilutions. Selected grids were loaded into a Titan Krios transmission electron microscope (FEI, ThermoFisher Scientific) operating at 300 keV at liquid nitrogen temperature. Micrographs (dose-fractioned movies) of all the samples were acquired using a Falcon 3EC Direct Electron Detector (FEI, ThermoFisher Scientific) in electron counting mode (50 frames; dose rate was 1 e^-^/Å^2^ per frame) with a calibrated pixel size of 0.832 Å, and a nominal defocus range of 0.5 to 3 μm. We used the semi-automated acquisition program EPU (FEI, ThermoFisher Scientific) for data collection during several days of the eight different datasets (Fig. S2A).

### Cryo-EM data processing

Comparative initial analysis of the data was performed with RELION 2.1.0-3.0 ^38, 77^ and cisTEM 1.0 ^78^. The final processing workflow was implemented in RELION and was applied in the same way to the 8 different datasets (Fig. S2). The individual frames of the micrographs were aligned and dose-weighted using MotionCor2 ^79^, and CFFIND-4.1 ^80^ was used to estimate the contrast transfer function (CTF) parameters of the aligned images without dose-weighting. The calculated parameters were used to discard bad micrographs. As a first approximation, manual picking was done in a small number of micrographs to optimize the parameters for automatic picking, designed to select both assembled and disassembled ribosomal particles. Particles were extracted using a box size of 500 pixels and subsequent 2D and 3D classifications allowed the isolation of the different types of particles (70S, 50S and 30S, Fig. S2B-C). 3D initial models were computed (RELION and cisTEM) independently for the three particles sizes; binned data was used in the intermediate steps to accelerate processing. As a first approach to obtain 3D reconstructions of these three categories of particles, 3D classifications and refinements were optimized to use as many good particles as possible, avoiding subclassification into different conformations (Fig. S2D, grey highlight I). At this stage we obtained good maps for the 70S (Fig. S2D, S3A-C, S3H-J) and 50S particles, but the 30S reconstructions were suffering from preferential orientation issues. These global refinements of the 70S particles showed poor and diffuse densities for their 30S region (Fig. S3A, S3H), so in order to improve the quality of the maps without sacrificing the number of particles, a second branch of the processing workflow was performed in which the local refinements of the 50S, 30S-head and 30S body were calculated with the use of selective masking and signal subtraction (Fig. S2D, green highlight II; Fig. S3D-E, K-L). The improved maps were used for building and refinement of two initial models, R and L, that were utilized as templates for all the subsequent characterizations (Fig. S3F-G, M-N). A final fork of the processing workflow was performed to analyze the heterogeneity of the 70S particles in all datasets (Fig. S2D, pink highlight III). For that purpose, we used cryoDRGN 1.0 ^40^, a generative network-based algorithm that was able to approximate the heterogeneous nature of the 70S particles to a continuous function describing the different conformations of the ribosomal samples and their associated particles. To optimize the training of the model, the disassembled particles were excluded from the computation and, after testing different settings, the particle images were downsampled (3.25 Å/pixel) and a latent dimension of 8 with a large architecture (3 hidden layers with 1024 nodes each, for both the encoder and the decoder networks) was used for 50 epochs. Results were analyzed using cryoDRGN python-based tools and 10 classes per dataset were obtained after GMM clustering (Gaussian mixture model, Fig. S2D, III). The number of clusters was decided trying to find a balance between reducing the heterogeneity and keeping enough particles per class. Particles from these clusters were extracted and further processed in RELION for additional validation, analysis and regrouping of some classes. Mutant datasets seem to be more heterogeneous than the wild-type ones (7-9 against 10-11 classes) and regardless of the differences, the conformational landscape captured in cryoDRGN latent space is consistent between the datasets, despite being independently processed (Fig. 3A, 4D, S2D, S4-5). Final 3D refinements of the different classes were computed with RELION, with maps ranging most of them between 2.7 and 3.5 Å (2 halves gold-standard resolution estimation, Fig S4-5 and Data S2). All maps were sharpened by applying a negative B-factor, estimated by the post-processing method in RELION, and corrected for the modulation transfer function of the detector. Local resolution estimations and resolution-filtered maps were also obtained within RELION (Fig. S4-5). Additional post-processed maps were obtained using Phenix^81^ and LAFTER ^82^.

### Model building and structure refinement

Initial models for the PAO1 ribosomal proteins were obtained with the use of the structure homology-modelling tools Phyre2 ^83^ and Swiss-Model ^84^. The initial model of the ribosomal RNAs was derived from *E. coli* ribosome structure (pdb: 4YBB). Initial fitting and analysis were performed in ChimeraX^85^ and real-space refinement were performed using Phenix ^81^. COOT^86^ was used for model building and modification, manual local refinement and visualization. UCSF ChimeraX was also used for model and map inspection, structure modification and map segmentation and modification. Figures including map densities, local resolution and coordinates were generated in Chimera/ChimeraX or PyMOL (Schrödinger, L., & DeLano, W. (2020). *PyMOL*. Retrieved from http://www.pymol.org/pymol).

### Structural analysis

Once we obtained from the 8 datasets their 78 final maps and models (Fig. S4-5) we try to characterize in conformational terms the associated variability of the system, the underlying complexity described in the latent spaces computed by cryoDRGN. As a first simple approximation, we decided to calculate the distance between the structural elements of the 30S body and the 50S subunit (Fig. S6A-B). This is clearly an oversimplification, as for example similar distances can be obtained in different ways, but it provides a first helpful glimpse about how the system behaves when analyzing the open-closed state of the ribosome. The different conformations of all datasets (ribosome + antibiotic) were organized according to these distances and their relative contribution to the total population calculated and analyzed (Fig. 3A). This structural classification allowed us to highlight common features related to the changes produced by the addition of antibiotics or the presence of the uL6 mutation, and the movement transitions between the different conformations inside the datasets. Distances (here or in subsequent analysis; defined between Cα or P atoms) were calculated either inside Chimera or by extracting data directly from the PDB files using Python scripts. After getting this basic description of the heterogeneity and transitions of the system, we decided to obtain a deeper and unifying description of the whole system. In this approach, the variability is described in terms of the movement of the different rigid bodies that compose the ribosome ^58^. Following a first analysis of the changes happening in the different conformations, we decided to divide the ribosome into the next rigid bodies (Fig 4A): head (16S (917-1382 nt), S3, S7, S9, S10, S13, S14, S19), B1 (16S (1-559 nt, 589-634 nt, 1416-1471 nt) S4, S12, S16, S17, S20) and B2 (16S (560-588 nt, 635-916 nt, 1384-1415 nt, 1472-1527 nt), S5, S6, S8, S11, S15, S18, S21) domains of the 30S subunit, and the 50S subunit (23S, 5S, L2, L3, L4, L5, L6, L9, L13, L14, L15, L16, L17, L18, L19, L20, L21, L22, L23, L24, L25, L27, L28, L29, L30, L32, L33, L34, L35, L36). To describe changes between conformations, displacement vectors were visualized in Chimera using its ability to interpret bild files (containing the Cα/Phosphate coordinates of the initial and final models that describe the transition; Fig. 4B). In addition, rotation axes were calculated with the structure analysis tools in Chimera to summarize the movements between two conformations in a transition and the changes in the associated contact surfaces (Fig. 4B-C).

We decided to take the Ro1 conformation as a reference. This is due to two reasons, i) it is the most abundant class of the wild type without antibiotic dataset, and ii) it is at the opposite end of the transition that characterizes the changes induced by the presence of the AGAs or the mutation (Fig. 3A). Rotation axes (of the 50S, B1 and 30S-head) were calculated for the transitions between Ro1 and all the other classes, with all the structures fixed in the described B2 domain (containing the DC and the AGA binding sites; RMSD B2fit = 0.244-0.961 Å). The main purpose of this method is to achieve a systematic comparison of all the conformations (Fig. 4D). This can be also interpreted also as the possible movements needed to go from the open conformation Ro1 to each of the other conformations in the datasets. We found that we could summarize the conformational landscape of the datasets by plotting only the values of the rotation of the 50S and B1 domain (Fig. 4D). This oversimplification removes additional data, like the different directions of the rotational axes and their associated small translations, but allows us to correlate different datasets and classes, and easily characterize diverse clusters in the data that rationalize the original latent spaces used to obtain the classes. In addition, this approach explains the previous D values as the result of two concurrent independent rotations, with the particularity that the extension of the changes depends on the distance to the rotation axis (Fig. 3A, 4C-D).

Within this framework, we defined different clusters that can be used to describe the transitions induced by the addition of antibiotics and the presence of the uL6 mutation (see main text). As previously mentioned, there are also changes in the direction of the rotation axes and cluster (a) can be used to show an example of it (Fig. S7A). We can identify two groups of rotations: LTc1, LTc3 and LTc6 show just small deviations, while LTc2, LTc5 and LTc7 display larger changes when compared with the previous group. All these variations are the result of different interactions between the domains, compositions (the presence of tRNAs) and configurations (for example, rt *vs* st). In this respect, the presence or absence of interaction between the 50S and the 30S-head domains affects the rotation axis of the 50S (Fig. S7A right panel). When comparing members of different clusters (Fig. S7B), for example, LTc1 (a) and RTc2 (b), we can see that the differences, in this case, are mainly due to the values of the B1 rotation, while if we compare both with RTc7 (c) the differences are related both with the change in the direction and the magnitude of the rotations. In this regard, cluster (e) is characterized by larger rotations of the 50S, and variations in both the direction and value of the B1 rotation that can be interpreted as counterweights of the former (Fig. S7C). Cluster (f) is additionally described by the release of the 30S-head from its interaction with the 50S subunit, which contributes with the instability of these classes as additional intersubunit bridges are lost (Fig. S7D). This effect can be identified as a pure rotation of the 50S departing from the 30S-head, with a rotation axis that is sort of perpendicular to the previous examples (Fig. S7D top panel). In other cases, it is mixed with a more typical 50S rotation, producing a deviated rotation axis (Fig. S7D bottom panel).

The identification of classes with singular composition helped in the interpretation of the data. In this regard, we defined AP(E) classes as those containing densities corresponding with tRNAs and mRNA. Due to the compositional heterogeneity of the purified samples, those densities are not well defined, especially in the case of the E position (Fig. S6C). We were also able to detect SmpB-tmRNA complexes (Fig. 5F). Interestingly, while the AP(E) complexes were apparently disassembled by the presence of tobramycin, the tmRNA complexes were resistant to it. The AP(E) classes were used to make a focused comparison of the datasets, as they represent the same composition (Fig. 5A-E). For that purpose, in addition to the previous rotational analysis (Fig. 5A-D) we calculated the distance between equivalent residues (Cα and P coordinates) of Rc2 (reference, as it is the wild-type AP(E) class without antibiotics) and the rest of AP(E) classes (all fitted in their B2 domain; Fig. 5E). To extend this interpretation of the data, as changing accessible and intersubunit contact surfaces, we analyzed also other conformations (a tmRNA class and main representative classes of the R, L, LT, LK and LC datasets; Fig. S7E). The especial composition of the tmRNA class LTc5 produce larger changes in the 50S subunit and the 30S-head than for example LTc6, although the B1 changes look similar (Fig. S7E, 5E, 4D). The Ro1 class experiences changes in the opposite directions, as it is happening also with RCi3 and LCo2. Lc3, LTc1 and LKc3 show equivalent changes when compared with their AP(E) counterparts, but with a common larger rotation of the head (and different to the LTc5 one; Fig S7E, 5E).

Unexpectedly, a weak extra density can be recognized in some of the classes with free 30S- heads, both R and L (Fig. S7F). Merging some of these classes, reclassifying and performing local refinements of the 30S-head allowed us to isolate the density and identify it as lysozyme contamination coming from the lysis buffer, with the help of mass spectrometry analysis. It was first identified as some kind of translation factor, because of its specific position between the head and the body of the 30S subunit (Fig. S7F), similar to the location of the HPF ^87^. Its faint density indicates that its occupancy is low, and it is always associated with a flexible 30S-head that must separate quite enough from the B2. Despite its nearness to h44 there is no correlation between its presence and lower occupancy of AGAs, but as it stabilizes the release of the 30S-head from the 50S subunit a role in the ribosomal disassembly cannot be discarded. Although this is a purification artifact, previously described ^88^, it is tempting to suggest that maybe this interaction could conduce to the sequestration and inactivation of ribosomes from disrupted bacteria in the context of an infection, inside or outside phagocytes (lysozyme is one of the markers of mononuclear phagocytes ^89^, maybe preventing the uncontrolled synthesis of toxins or random antigens. This would provide additional support for the use of lysozyme as an alternative antibiotic^90^ ^91^ and the described interaction could constitute the base for the design of new drugs.

## SUPPLEMENTARY FIGURE LEGENDS

**Fig. S1 Minimum inhibitory concentrations (MICs) for various antibiotics and sequence alignment. A)** Tobramycin, Tetracycline, Carbenicillin, Ceftazidime, Erythromycin, Ciprofloxacin, Colistin and Aztreonam antibiotics MIC were tested by analyzing the maximal growth after 48h in the presence of each antibiotic. **B)** Alignments of 3310 uL6 protein sequences from *P. aeruginosa*. Only sequences with alterations compared to PAO1 in the highly conserved region of uL6 are shown. The asterisk denotes the reference laboratory strains PAO1 and PA14. In bold the mutation in our collection of clinical strains of *P. aeruginosa*.

**Fig. S2. Single-particle cryo-EM of R and L ribosomal samples. A)** Typical cryo-EM micrographs of R and L ribosomal particles in vitreous ice imaged with a Titan Krios Falcon 3 direct detector camera (the white scale bars included in the images represent a length of 200 Å). Particles are more heterogenous in size at the right panel (L) due to the presence of more disassembled particles. **B)** Characteristic 2D averages of the different particles present in the partially disassembled mutant ribosomal sample. **C)** Representative reference-free 2D class averages of R 70S mutant ribosomal particles computed by RELION. **D)** Comparative cryo-EM data processing workflow of two of the datasets (out of eight): R (blue) and L (red), without antibiotics. Each final 3D map is represented by its name, number of particles and global resolution. After initial preprocessing of the data, three types of 3D reconstructions were obtained with three differentiated goals: (I) Global refinements, performed with as many particles as possible, describing the particles and their heterogeneity (in the case of the 70S particles, fuzzy 30S subunits); (II) Local refinements, where masking and signal subtraction were used to locally refine the reconstruction of the 50S, the 30S-body and the 30S-head (for analysis and model building, see Fig. S3); (III) Homogeneous refinement of discrete clusters of particles calculated with CryoDRGN. UMAP dimensional reduction plots (top-left panels) show the distribution of particles in the latent space. Clustering of the particles (top-right panels) provided initial classes (bottom panels) that, after analysis, cleaning, and re- classification, were used to produce the final homogeneous refinements representing different conformational states of the particles (see Fig. S4-5, and Methods section for details).

**Fig. S3. Resolution assessment and validation of the final cryo-EM reconstructions. A-G**) Analysis of the globally and locally refined R maps. **H-N**) Analysis of the L maps. **A, H)** Final global reconstructions filtered and colored according to the local resolution estimated by RELION. **B, I)** Angular distribution of the particles used for calculating the final global 3D reconstructions (A/H). **C, J)** Directional FSC plots for the reconstructions calculated with 3DFSC ^92^. Sphericity, as a measure of the low anisotropy of these reconstructions, is indicated above the graphs. **D, K)** Two different views of the synthetic maps generated from the local refinements of the 50S, 30S-body and 30S-head regions (shaded at the small insets in blue, yellow and red respectively). Maps are colored according to the local resolution, with the same gradient key as at A/H and show a considerable improvement in the details of the 30S subunit. **E, L)** Cutaway views of the synthetic maps colored with a more focused range of resolution values. **F, M)** Models derived from the local 3D reconstructions. **G, N)** Fourier Shell Correlations (FSC) curves of the final maps and local reconstructions, with global resolution values estimated according to the gold-standard criterion (FSC=0.143).

**Fig. S4. Local resolution and resolution assessment of the 35 cryo-EM R maps presented in this work.** Each map was post-processed using RELION for the local resolution estimation and local filtering. A common color key is used for all the maps (see in A). **A)** R 50S disassembled map. **B)** Wild-type ribosomal maps without antibiotics. **C, E, G, I)** FSC curves of the preceding group of maps, used to calculate their global resolution values according to the gold-standard criterion (FSC=0.143). **D)** R maps in the presence of Tobramycin. **F)** R reconstructions with Kanamycin. **H)** Maps of wild-type ribosomes incubated with Chloramphenicol. The name of the characteristic conformation of each reconstruction is indicated by its global resolution, local resolution range (at the visualized density level) and the number of particles used in the final 3D refinement.

**Fig. S5. Local resolution and resolution assessment of the 45 cryo-EM L maps presented in this work.** Each map was post-processed using RELION for the local resolution estimation and local filtering. A common color key is used for all the maps (see in A). **A)** L 50S disassembled map. **B)** Mutant ribosomal maps without antibiotics. **C, E, G, I)** FSC curves of the preceding group of maps, used to calculate their global resolution values according to the gold-standard criterion (FSC=0.143). **D)** L maps in the presence of Tobramycin. **F)** L reconstructions with Kanamycin. **H)** Maps of mutant ribosomes incubated with Chloramphenicol. The name of the characteristic conformation of each reconstruction is indicated by its global resolution, local resolution range (at the visualized density level) and the number of particles used in the final 3D refinement.

**Fig. S6. Characterization of structural features in R and L classes. A)** Definition of the distance D between nucleotides A55 of helix h15 (30S) and G2649 of the SRL (50S), chosen to characterize the conformational state of the R and L cryo-EM maps. The main panel shows two examples of these distances (Ro1, open and blue; LTc1, closed and orange). Both structures were fitted in their 50S subunit for comparison purposes, so the open or closed state of the 30S-body can be illustrated. **B)** The closing mechanism of the 30S subunit is associated with the function of EF-Tu in translation. The right panel depicts the interaction of EF-Tu with the helix h15 of the 30S subunit. **C)** Representation of the densities associated with tRNAs in the AP(E) classes. **D)** View of the Chloramphenicol binding site in maps RCci7 and LCo2. **E)** Depiction of the 50S vulnerable region, H69 and uL6 densities in the two rt classes of the mutant in the presence of Chloramphenicol (see Fig. 2G for details and comparison).

**Fig. S7. Structural analysis of the identified clusters. A)** Comparison of the rotation axes of some members of cluster (a) in 4C. Three perpendicular views are shown, as in 4B. The displacement vectors correspond with the transition Ro1 to LTc7. The right panel illustrates the orientation of the 30S-head on three of the classes shown on the left panels. **B)** Comparison of the rotation axes of members of clusters (a), (b) and (c). The displacement vectors display the Ro1 to RTc7 transition. **C)** Characteristic displacement vectors (Ro1 to RTt1) and rotation axes of cluster (e) (tight conformations). **D)** Effect of the release of the 30S-head on the orientation of the 50S rotation axes of two classes. **E)** Additional analysis of the changes in the accessible surface and the intersubunit interface of ribosomal conformations (see Fig. 5E). **F)** Identification of the weak signal in certain classes as lysozyme. The top left panel shows the presence of low densities (yellow) between the head and the body of the 30S subunit of class Lo1. Its displacement vectors are shown at the top right panel, supporting the large rotation of the 30S to accommodate the lysozyme monomer. Detailed views of the lysozyme density are shown at the bottom panels, indicating the interacting helices and nucleotides.

**Fig. S8. Additional characterization of the Tobramycin binding sites. A)** View of the relative location of the five Tobramycin sites in a slice of the ribosome (see lower inset). **B)** Occupancy gradient for Tobramycin site 3. **C)** Depiction of the absence of densities for Tobramycin in the control classes incubated with Chloramphenicol. **D)** Local changes associated with the binding of Tobramycin into the five sites.

**Data S1 Proteomics comparison between PAO1 and L6 mutant strains.** Comparative proteomics analysis of the whole cell proteome of the L6M mutant strain relative to the wild type PAO1. Only proteins with at least a 1.5-fold change relative to the wild-type strains (p > 0.05, Anova) and with 3 identified unique peptides were reported as differentially expressed. (see Excel table DataS1)

**Data S2 cryo-EM data collection and refinement table.** (See Excel table DataS2)

